# A genetically encoded RhoG FRET biosensor reveals spatially compartmentalized RhoG–Rac1 signaling during cell protrusion

**DOI:** 10.64898/2026.08.24.746776

**Authors:** Maíra de Assis Lima, Abigail Thomas, Roshan Ravishankar, Rafael Garcia-Mata, Gaudenz Danuser, Veronika Miskolci, Dianne Cox, Louis Hodgson

## Abstract

RhoG is a member of the Rho-family of small GTPases, and is closely related to the canonical Rac1 GTPase, implicated in membrane trafficking, dorsal ruffling, macropinocytosis, and cell protrusion, but its activity has been difficult to visualize directly in living cells with high spatial and temporal resolution. Here, we developed and validated a genetically encoded, single-chain Förster resonance energy transfer (FRET) biosensor for RhoG based on a C-terminal full-length RhoG and an intramolecular RhoG-binding domain derived from ELMO1. The biosensor showed a robust dynamic range when comparing constitutively active and inactive RhoG mutants, responded appropriately to regulation by RhoGDI, GAPs, and GEFs, and detected growth factor–stimulated RhoG activation in live cells. Imaging in mouse embryonic fibroblasts revealed dynamic RhoG activation at leading-edge protrusions, dorsal ruffles, and forming pinocytic and macropinocytic structures. To define the signaling relationship between RhoG and its closely related family member Rac1, we combined the RhoG biosensor with a near-infrared Rac1 FRET biosensor for simultaneous live-cell imaging. Morphodynamic mapping showed that both RhoG and Rac1 activities were positively coupled to edge protrusion, with strongest correlations near the leading-edge, but their direct coupling varied with distance from the edge, indicating partial spatial decoupling within protrusive regions. Inhibition of Src-family kinases altered RhoG dynamics, strongly suppressed Rac1 coupling to protrusion, and inverted the normal positive correlation between RhoG and Rac1 activities. Signaling microdomain analysis further showed that Src inhibition selectively prolonged Rac1 microdomain lifetimes without significantly affecting RhoG domains. Together, these results establish a new biosensor for direct visualization of RhoG activity and reveal that RhoG and Rac1 are coordinated but spatially and temporally distinct components of protrusion-associated signaling networks, with Src-family kinases playing a central role in maintaining their normal coupling.

## Introduction

RhoG is a member of the Rho family of p21 small GTPases and is most closely related to Rac1 (1–5). Its expression patterns are widespread with relatively low tissue specificity, but are associated most highly with cells of hematopoietic origin, including monocytes and neutrophils, as well as lymphoid tissue and bone marrow, skin and brain (6–9). Importantly, RhoG has been associated with cell motility regulation, both in normal physiology and in diseases including cancer (2, 10–12). Like other Rho-family GTPases, RhoG functions as a molecular switch that cycles between an active, GTP-bound state and an inactive, GDP-bound state. Although historically less studied than the canonical Rho GTPases, increasing evidence indicates that RhoG plays important roles in cytoskeletal remodeling, cell migration, dorsal ruffling, macropinocytosis, and phagocytosis, particularly in immune and epithelial cells (1, 2, 13–28). RhoG exerts many of these functions through signaling pathways that engage ELMO–Dock complexes and through functional crosstalk with Rac1 (13, 16, 29, 30). These properties place RhoG within a broader signaling network that coordinates membrane dynamics, actin remodeling, and cell motility. Despite its relevance to these processes, tools to directly visualize the spatiotemporal regulation of RhoG in living cells have been lacking.

Cell protrusion is driven by highly localized and rapidly changing signaling events at the plasma membrane, where Rho-family GTPases organize actin polymerization, membrane deformation, and adhesion-associated signaling (31–36). Previous work has shown that related GTPases such as Rac1, Cdc42, and RhoA display distinct spatial and temporal relationships to protrusion and retraction cycles, indicating that their signaling is tightly patterned within the leading-edge rather than uniformly distributed throughout the cell (37–44). RhoG has been implicated upstream of Rac1 activation and linked to membrane ruffling and endocytic structures (13, 15, 16, 27, 45–47), and participates in signaling programs during protrusive behavior (48). However, without a direct activity reporter, it has remained difficult to determine where and when RhoG is activated during edge dynamics, how closely its activity is coupled to protrusion, and to what extent its signaling is coordinated with other GTPases, including Rac1 in living cells.

In addition to large-scale edge dynamics, cell motility is increasingly understood to depend on small, spatially restricted signaling assemblies that form, persist, and turnover within subcellular regions (49–51). Such local signaling clusters, or signaling microdomains (MD), are thought to provide a mechanism for organizing signaling specificity in space and time, allowing GTPase activities to be coordinated within localized membrane or cytoskeletal environments rather than broadly across an entire cell (52). For Rho-family GTPases, this type of local organization may be particularly important at protrusions, dorsal ruffles, and trafficking-associated membrane structures, where signaling must be both dynamic and spatially constrained. Whether RhoG activity is organized into such local MDs, how these domains relate to other MDs, including those of Rac1, and how upstream kinases influence their persistence and coordination has yet to be studied.

Genetically encoded, single-chain Förster resonance energy transfer (FRET) biosensors provide a powerful and a user-friendly approach to address these questions by enabling direct visualization of protein activity in living cells with high spatial and temporal resolution (40, 42, 44, 53). These biosensors typically consist of a donor and acceptor fluorophore connected to a Rho GTPase and a GTPase-binding domain derived from a downstream effector (54, 55). Upon activation of the GTPase, an intramolecular conformational change alters the relative orientation of the fluorophores and modulates FRET coupling. Unlike other intermolecular or bimolecular designs for Rho family GTPases that have been available in the field (39, 56–58), the single-chain design ensures equimolar distribution of the FRET donor and acceptor moieties everywhere within a cell, making for simple and straightforward image acquisition and data processing pipelines. This design enables reversible, ratiometric imaging of GTPase activity in intact cells and has been successfully applied to Rac1, RhoA, Cdc42, and others during migration, polarity establishment, and membrane protrusion (38, 59–66). Extending this approach to RhoG would provide an opportunity to define its signaling behavior readily and directly, distinguish its dynamics from those of closely related GTPases, and establish how it contributes to protrusion-associated signaling networks.

Here, we report the development of a new genetically encoded, single-chain FRET biosensor for RhoG. The biosensor uses circularly permuted monomeric ECFP and monomeric Citrine as the donor and acceptor fluorophores and is designed to preserve the full-length RhoG sequence, including its C-terminal hypervariable region and the prenylation motif, to maintain proper membrane targeting and interaction with upstream regulators. We first characterized the biosensor and then applied it in mouse fibroblasts to visualize RhoG activity during growth factor stimulation and spontaneous protrusive behavior. To define the relationship between RhoG and Rac1 in cell morphodynamics, we combined this biosensor with our previously developed near-infrared (NIR) FRET biosensor for Rac1 (67), enabling simultaneous imaging of the two GTPases in the same living cells. Using morphodynamic mapping (39), we analyzed the spatial and temporal coupling of RhoG and Rac1 activities to constitutive protrusion–retraction cycles and examined how Src-family kinases influence this coordination. We further used signaling MD analysis (68) to quantify the organization, persistence, and colocalization of RhoG- and Rac1-associated activity clusters. The single-chain, genetically encoded biosensor for RhoG enables investigation of how RhoG contributes to protrusion, membrane remodeling, and localized signaling architecture in living cells.

## Results

We generated a genetically encoded, single-chain FRET biosensor for RhoG (Fig. 1A). The biosensor was based on a backbone originally derived from our Rac1 FRET biosensor (60), but incorporated a single affinity domain from ELMO1 (amino acids 1–115) (13) linked by a flexible linker of optimized length (69). The ELMO1-derived Rho-binding domain selectively pulled down activated RhoG mutants, including Q61L and G12V, but not the inactive mutant T17N (Fig. S1A,B). The biosensor contained the circularly permuted fluorescent proteins (70) mcp229ECFP and mcp157Citrine-YFP, each carrying the monomerizing A206K mutation (71). Full-length RhoG was fused at the C-terminus to preserve the hypervariable region and C-terminal lipid modification sites required for correct membrane targeting and interaction with RhoGDI (72–76). Detailed information on designing, engineering, and optimization of the single-chain RhoG FRET biosensor can be found in the Supplementary Text.

**Figure 1:**
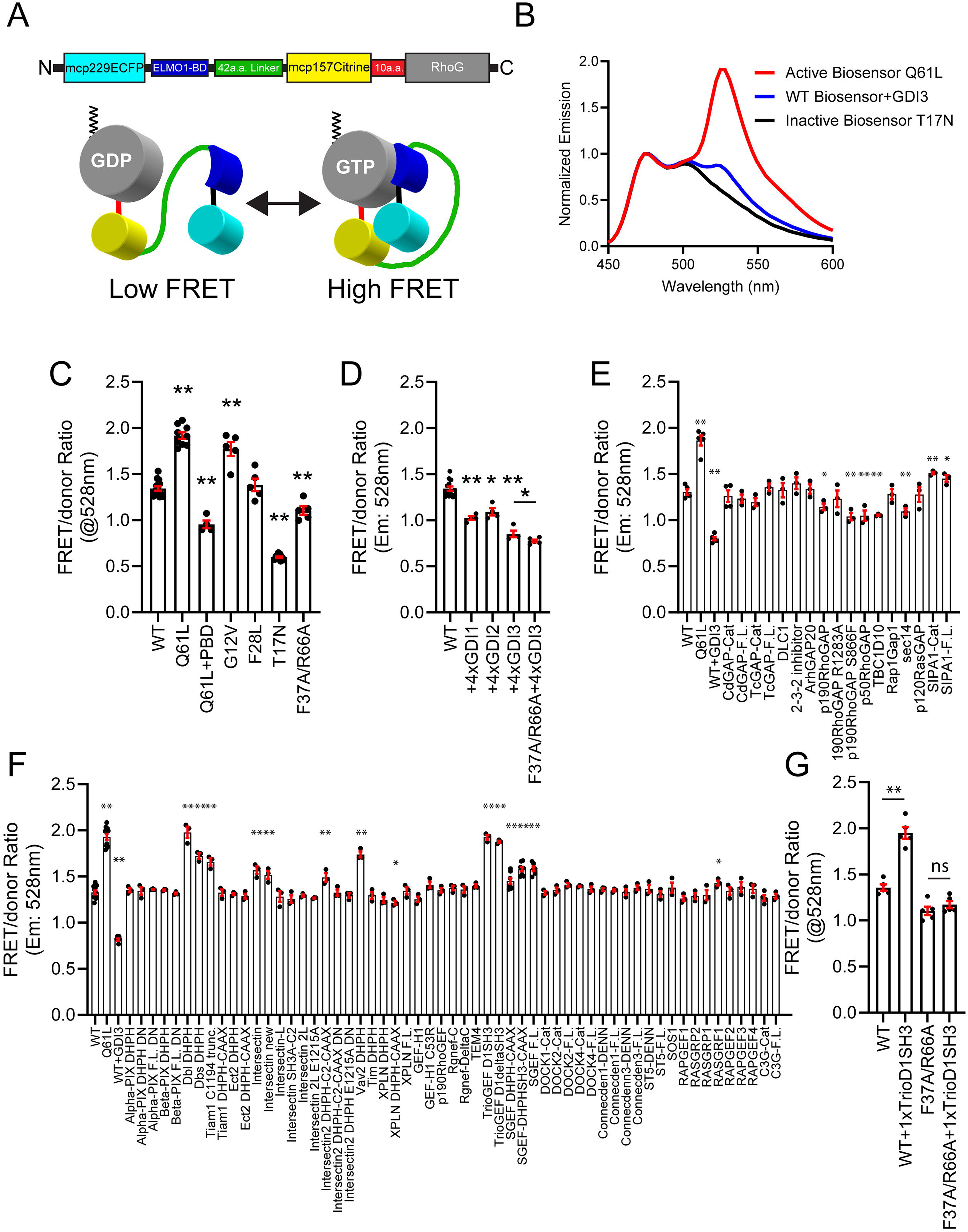
Development of a single-chain genetically encoded FRET biosensor for RhoG GTPase. A. Basic design of the RhoG GTPase FRET biosensor, based on cyan-yellow fluorescent proteins. The optimized biosensor consists of fluorescent proteins mcp229ECFP and mcp157Citrine-YFP. A full length human RhoG GTPase is attached at the C-terminus, via a 10 a.a. linker segment. An affinity binding domain derived from ELMO-1 is incorporated internally within the biosensor structure, followed by 42 a.a. segment of a flexible linker. B. Representative emission spectra following donor excitation in cells expressing constitutively active (Q61L) or dominant-negative (T17N) RhoG mutant biosensor. WT plus 4x excess GDI3 condition is also shown. C. Quantification of RhoG biosensor activity (FRET/donor emission ratio) measured in constructs carrying mutations affecting RhoG activation, effector, or the effector-binding interface, n=4-13. D. Quantification of RhoG biosensor response to co-expressed GDI isoforms. n=4-13. E. RhoG biosensor activity measured following expression of a panel of GTPase-activating proteins (GAPs), n=3-5. F. Quantification of RhoG biosensor response to co-expressed panel of GEFs, n=3-11. G. Quantification of RhoG biosensor response to TrioGEF catalytic domain co-expression, showing that competent effector binding interface is required for GEF-mediated activation of the biosensor, n=5. Data represent mean ± SEM. Statistical significance was determined using an unpaired two-tailed Student’s t-test. *p<0.05, **p<0.01.

Because the size of the biosensor precluded in vitro purification, we characterized it in HEK293 cells as previously described (38, 77). Wild-type (WT) and mutant versions of the biosensor were overexpressed in HEK293 cells, and fluorescence emission spectra between 450 and 600 nm were measured in detached cells following excitation at 433 nm. Constitutively active mutations in RhoG (Q61L and G12V) increased the FRET/donor ratio, whereas the dominant-negative mutation T17N reduced it. The difference in FRET/donor ratio between the active Q61L and inactive T17N biosensors was approximately 3.2-fold (Fig. 1B, C). In contrast, the active Q61L biosensor containing an unrelated affinity domain derived from p21-activated kinase 1 (PBD) showed a markedly lower FRET/donor ratio than the corresponding biosensor containing the competent ELMO1 RBD (Fig. 1C). This result indicates that the elevated FRET/donor ratio depends on specific intramolecular binding between activated RhoG and the ELMO1 RBD within the biosensor backbone. The fast-cycling RhoG mutant F28L, which does not require GEF for nucleotide exchange (78), did not increase the FRET/donor ratio relative to the activated mutants and instead behaved similarly to the WT biosensor (Fig. 1C), consistent with the GST-ELMO1 RBD pulldown results (Fig. S1A). In addition, introduction of the effector-binding mutations F37A (79) and R66A (29) into RhoG which are expected to impair binding to the ELMO1 RBD, reduced the FRET/donor ratio relative to the WT biosensor (Fig. 1C). Together, these results demonstrate that the FRET response of the biosensor is appropriately modulated by the activation state of RhoG and depends on its specific interaction with the cognate ELMO1 RBD.

We next tested regulation of the biosensor by RhoGDI isoforms (Fig. 1D). RhoGDI3 has been shown to interact with RhoG and to regulate its targeting to endomembrane and Golgi compartments, although broader RhoGDI isoform interactions may be context-dependent (80–82). RhoGDI1, RhoGDI2, and RhoGDI3 cDNAs were titrated against a constant amount of WT RhoG biosensor cDNA (Fig. S1C-E). All three isoforms reduced the FRET/donor ratio, but with distinct saturation profiles. As expected, RhoGDI3 produced the strongest reduction in FRET/donor ratio. The F37A/R66A effector-binding mutant remained responsive to RhoGDI3 and showed a slightly greater reduction in FRET compared to that of the WT biosensor (Fig. 1D), possibly favoring GDI interaction in absence of effector binding ability. We then screened for the response of the RhoG biosensor to co-expression of panels of GAPs and GEFs (Fig. 1E-G). Interestingly, co-expression of p50RhoGAP or p190RhoGAP reduced the FRET/donor ratio of the WT biosensor. For p190RhoGAP, the effect was observed with both WT p190RhoGAP and the activated S866F mutant, but not with the catalytically inactive R1283A mutant. These two GAPs have previously been shown to associate with the constitutively activated G12V RhoG (83, 84), but whether they serve physiologically relevant catalytic roles remain unknown. Expression of the RabGAP TBC1D10 also reduced the FRET/donor ratio, as well as Sec14, an inhibitor of the Dbs GEF (85), also reduced the FRET/donor ratio. Rap1GAP1 had no detectable effect, whereas SIPA1 produced a small but reproducible increase in the FRET/donor ratio. Among the GEFs tested, Dbs, Dbl (MCF2), Vav2, TrioGEF, and S-GEF increased the FRET/donor ratio, as expected (15, 46, 86–88). Interestingly, Tiam1 and Intersectins also increased the FRET/donor ratio which were unexpected and could be through indirect mechanisms by affecting other Rho GTPases, including Rac1 and Cdc42, and/or through interaction with other signaling complexes. RasGRF1 had only a marginal effect, whereas DOCK family GEFs and other Rab/Rap GEFs had no detectable effect, as expected. The biosensor carrying the F37A/R66A mutations did not respond to co-expression of the TrioGEF fragment that strongly increased the FRET/donor ratio of the WT biosensor (Fig. 1G), pointing to effector binding interaction being critical for increased FRET/donor of the biosensor when activated by competent GEF fragments.

We next evaluated the biosensor in mouse embryonic fibroblasts (MEFs) using live-cell imaging. Transient expression of the constitutively active Q61L and dominant-negative T17N biosensor mutants produced an approximately 30% difference in the whole-cell average FRET/donor ratio (Fig. 2A). Consistent with increased FRET, the Q61L biosensor showed a reduced donor fluorescence lifetime when transiently expressed in U2OS cells and analyzed by time-domain fluorescence lifetime imaging (Fig. 2B). To test whether the biosensor engaged endogenous downstream effectors, we performed a GST-ELMO1-RBD pull-down assay using the constitutively active Q61L biosensor containing either the ELMO1 affinity domain or a nonbinding PBD domain from p21-activated kinase 1. The activated biosensor was detected in the pull-down fraction only when it contained the nonbinding PBD domain (Fig. 2C), demonstrating preferential interaction of RhoG with the built-in ELMO1 binding domain. We then tested whether the biosensor responded to exogenous stimulation. Previously, Platelet derived growth factor (PDGF) stimulation has been shown to activate RhoG through the RhoGEF Trio in a Src- and PI3K-dependent manner during circular dorsal ruffle formation (27). In serum-starved MEFs, stimulation with PDGF (100 ng/mL) induced a time-dependent increase in the RhoG FRET/donor ratio. The response peaked at approximately 3 min and remained significantly elevated between 3 and 5 min after stimulation (Fig. 2D). Together, these results demonstrate that the biosensor is suitable for live-cell imaging, that changes in the measured FRET/donor ratio reflect corresponding changes in FRET state as confirmed by fluorescence lifetime imaging, that the response is driven by intramolecular binding rather than interactions with endogenous effector targets, and that the biosensor responds appropriately to extracellular agonist stimulation.

**Figure 2:**
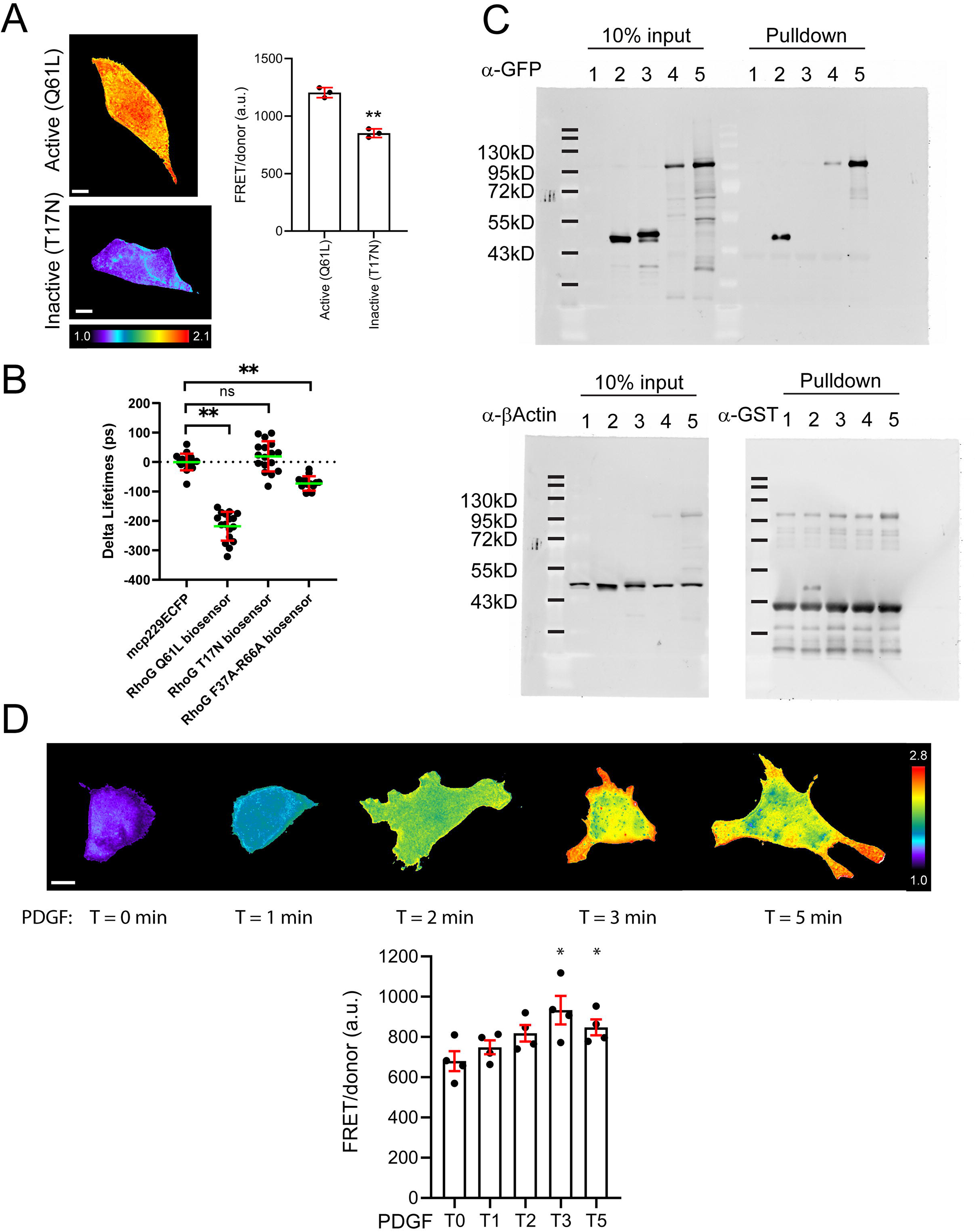
Characterization of a single-chain genetically encoded FRET biosensor for RhoG GTPase. A. Representative microscopy imaging-based analysis of activated (Q61L) versus inactive (T17N) mutant versions of the biosensor, overexpressed in MEFs (LEFT). White bars = 10 µm. Warmer pseudocolor ratio scale corresponds to higher RhoG activity. Quantification is also shown (RIGHT), n=3, shown with mean ± SEM. Statistical significance was determined using an unpaired two-tailed Student’s t-test. *p<0.05, **p<0.01. B. Time-domain fluorescence lifetime imaging analysis of FRET changes in activated (Q61L), inactive (T17N), and effector-binding mutant (F37A/R66A) versions of the biosensor transiently expressed in U2OS cells. Data are shown as changes in donor fluorescence lifetime relative to the donor-only control, which was used as the baseline for calculating FRET-dependent lifetime changes. Pooled data are from N = 3–4 independent experiments. Statistical significance was determined using an unpaired, two-tailed Student’s t-test. **p < 0.01; ns, not significant. C. Representative Western blot of the GST-ELMO-1 effector competitive pull-down analysis, showing preferential interaction of the cognate ELMO-1 binding domain with the activated RhoG within the biosensor. Lanes: 1, Untransfected control; 2, mGL-RhoG Q61L; 3, mGL-RhoG T17N; 4, biosensor containing RhoG Q61L; and 5, biosensor containing RhoG Q61L and ELMO-1 binding domain was replaced with PBD domain. D. Representative microscopy imaging-based analysis during exogenous agonist stimulation of starved MEFs using PDGF (100 ng/mL), showing images from t = 0 to 3 min time points (TOP). White bars = 10 µm. Warmer pseudocolor ratio scale corresponds to higher RhoG activity. Quantification is also shown (BOTTOM), n=4, shown with mean ± SEM. Statistical significance was determined using an unpaired two-tailed Student’s t-test. *p<0.05.

To examine subcellular RhoG activity patterns, MEFs expressing the biosensor were plated on fibronectin-coated coverslips and imaged during protrusion in serum (Fig. 3). RhoG activity was observed at leading-edge ruffling structures that subsequently collapsed (Fig. 3A; Supplementary Movie 1, 2). Following PDGF stimulation, edge ruffling was enhanced and frequently progressed into pinocytic vesicle formation (Fig. 3B, Supplementary Movie 3). During these events, RhoG activity first appeared as a central punctate signal within the forming pinocytic structure and then became more prominent over time throughout the vesicle (Fig. 3B). Dynamic RhoG activation patterns were also observed during dorsal macropinocytosis following PDGF stimulation (Fig. S2, Supplementary Movie 4).

**Figure 3:**
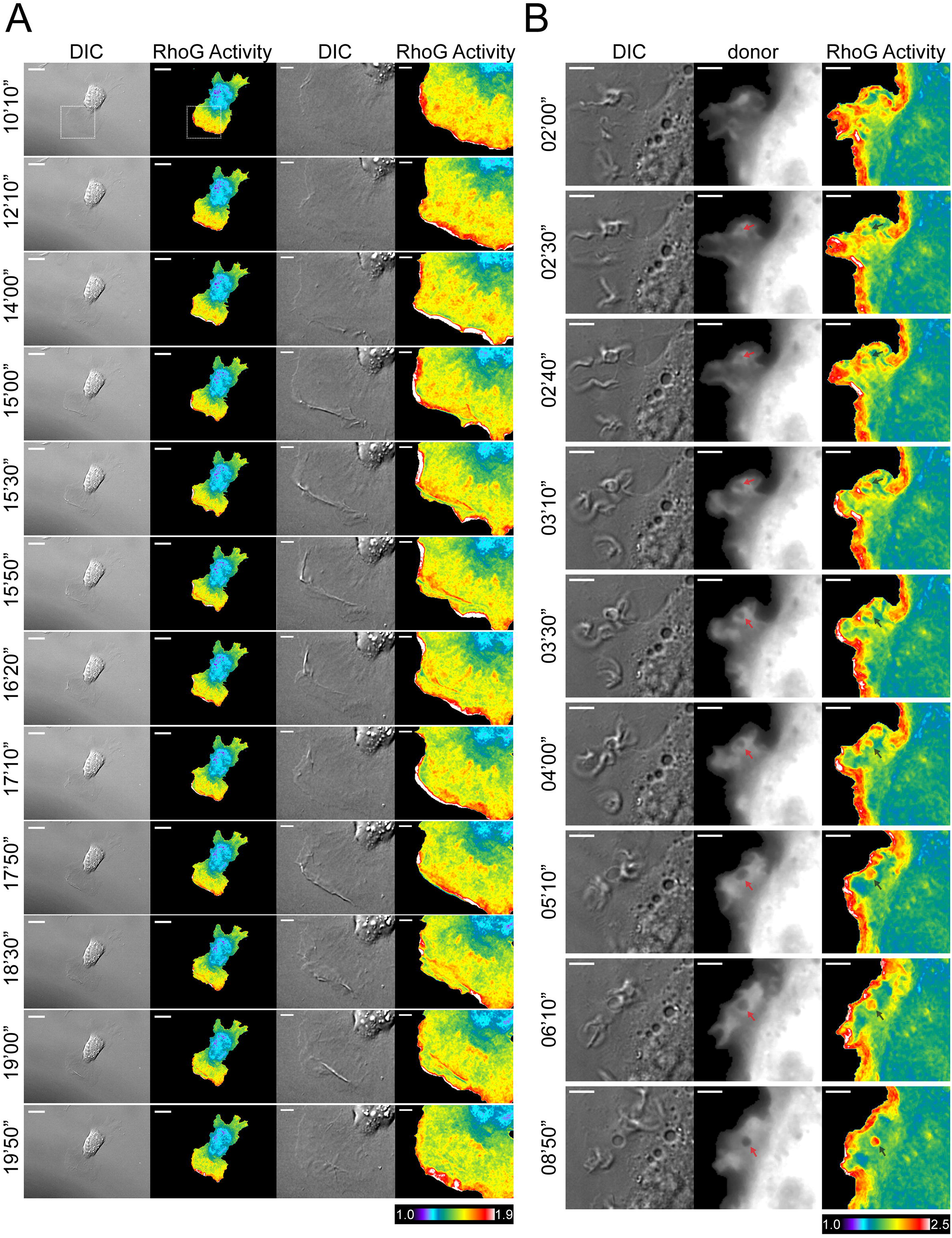
Representative, timelapse panels of RhoG activities in MEFs. A. Edge protrusion and associated ruffling in MEFs showing dynamic RhoG activities patterns during cell movement. White bar in the whole-cell image panels = 20 µm. Zoomed panels of the edge protrusion and ruffling are also shown. White bars = 5 µm. Warmer pseudocolor ratio scale corresponds to higher RhoG activity. B. Edge-associated pinocytosis in MEFs under PDGF-stimulated condition. Arrows indicate a forming pinocytosis structure. White bars = 5 µm.

Within the membrane protrusions, RhoG and Rac1 are known to be closely linked signaling nodes with a relatively well-established mechanistic connection, wherein RhoG can activate Rac1 through formation of ELMO1-DOCK complex, or the two pathways could operate in parallel (13, 16, 20, 27, 45, 46, 89–91). Thus, we sought to visualize RhoG and Rac1 activities simultaneously in living cells. To compare RhoG and Rac1 signaling dynamics in the same cells, we generated MEFs stably expressing the RhoG biosensor under a tetracycline-inducible promoter and then introduced our previously described near-infrared (NIR) Rac1 FRET biosensor, also under tetracycline control (67). Viral transduction, FACS sorting, and doxycycline concentration were optimized to achieve expression levels of approximately 34% of endogenous RhoG for the RhoG biosensor and approximately 21% of endogenous Rac1 for the Rac1 biosensor (Fig. S3), to achieve sufficient signal to noise ratio during live cell timelapse imaging. Expression of the biosensors did not alter endogenous RhoG or Rac1 levels (Fig. S3). We then monitored leading-edge protrusion–retraction dynamics in serum and analyzed the data using the morphodynamics mapping approach (39). Briefly, morphodynamics analysis tracks leading-edge motion and constructs dynamic measurement windows along the cell edge. These windows are then shifted rearward by defined distances to sample successive regions behind the edge during protrusion–retraction cycles. Within these spatially constrained windows, whose dimensions are selected based on the time resolution of the timelapse imaging sequence and diffusion-limited signaling dynamics, both edge protrusion velocity and protein activities, such as RhoG and Rac1, can be measured and cross-correlated across space and time, either with each other or with the edge protrusive velocity (39). This analysis enables visualization and quantification of protrusion–retraction cycling and spatiotemporal cross-correlation between cell edge motion and signaling activation at the leading edge and in regions behind the edge (Fig. 4A). Under control conditions, RhoG activity showed positive cross-correlation against protrusion velocity, with the strongest peak occurring at zero time-lag (Fig. 4B, Fig S4A) indicating direct concurrence of RhoG activity cycling with the leading-edge protrusive motion. Although coupling at the extreme edge was weaker, the maximal positive cross-correlation was detected at positions 3–6 pixels (0.9 – 1.8 µm; each pixel corresponds to approximately 300 nm) behind the edge, while maintaining zero time-lag against the edge protrusive velocity. The coupling of activity to protrusion then declined in strengths at greater distances from the edge. These observations indicated RhoG to be a strong and direct signaling node that is concurrently coupled to edge movement. We then turned to the analysis of Rac1 activity in the same cells. Under control conditions, Rac1 also showed positive cross-correlation with protrusion velocity (Fig. 4C. Fig S4B). The maximal cross-correlation occurred at 3 – 6 pixels (0.9 – 1.8 µm) behind the edge, similar to RhoG, and then decreased farther from the edge. The peak time-lag for Rac1 was slightly positive to suggest a leading function in controlling the cell edge protrusive initiation, although the 95% confidence interval spanned zero time-lag to indicate that it is actually indistinguishable in time from a direct concurrence, similar to RhoG (Fig. S5A, B). We then directly compared the two biosensor activities without referencing the protrusive edge velocities, as each measurement window enabled simultaneous recoding of both RhoG and Rac1 activities. The direct comparison of the RhoG and Rac1 biosensor signals showed strong positive cross-correlation at 0 – 3 pixels (0 – 0.9 µm) from the edge with zero time-lag (Fig. 4D), indicating a direct concurrence with strong coupling between these two signaling nodes. Interestingly, the coupling strengths decreased to nonsignificant levels between 3 and 15 pixels (0.9 – 4.5 µm) from the edge and then strongly increased again farther from the edge to approach similar levels to the cross-correlation coefficient peak value at the edge of the cell (Fig. S4C). These observations point to a highly coordinated but non-homogeneous and spatially distinct signaling compartments that exist within cell edge protrusions for RhoG and Rac1.

**Figure 4:**
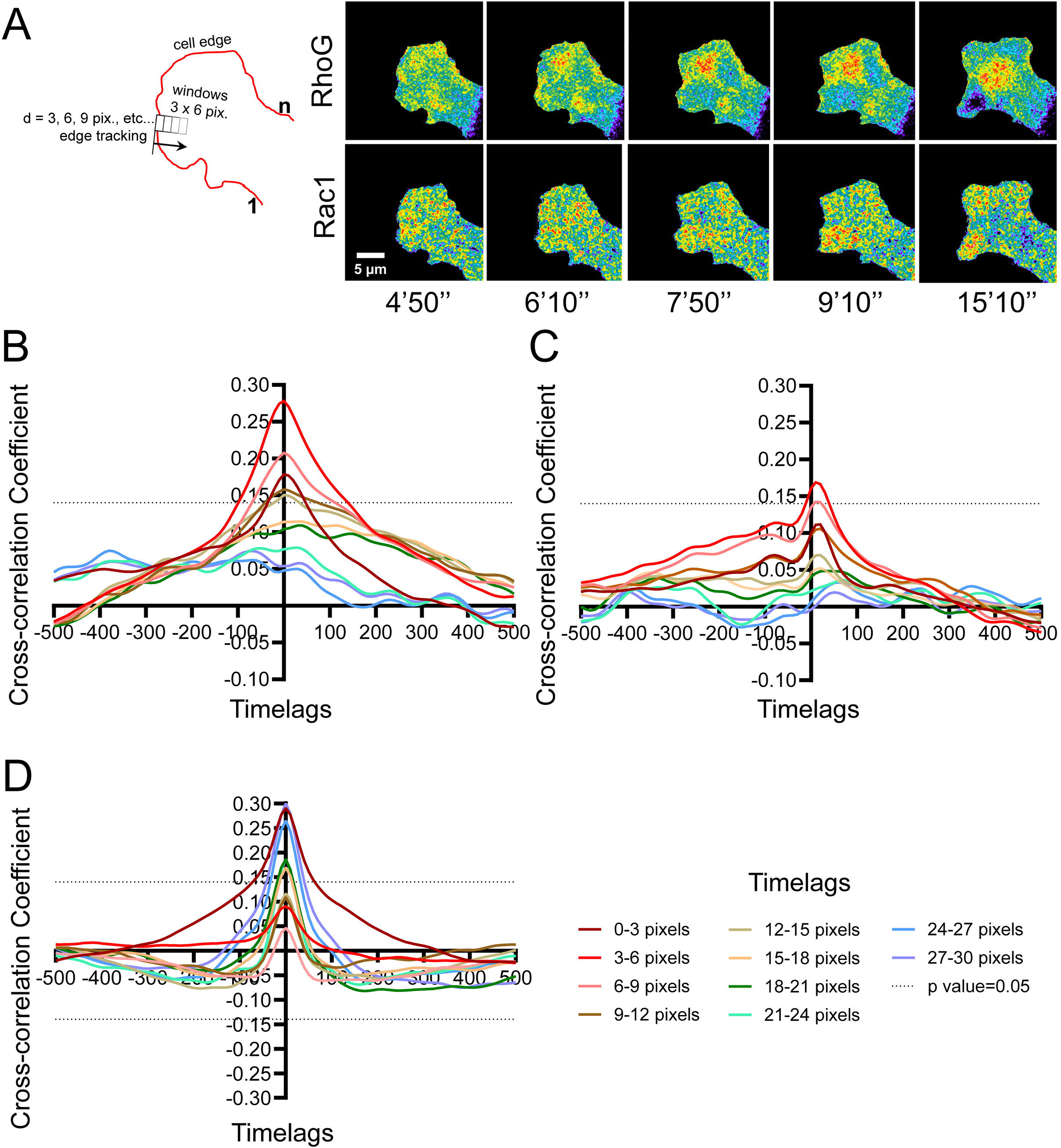
Morphodynamics mapping and cross-correlation analysis between RhoG and Rac1 activities within the leading-edge protrusions in MEF, as distances away from the edge. A. Schematic diagram of the leading edge measurement window construction and propagation in space as a function of time. Also shown are representative timelapse panels of RhoG and Rac1 biosensor activities in the same cell at the leading edge. White bar = 5 µm. Pseudocolor range: 1.0 – 1.3. B. Cross-correlation functions of RhoG activity against the leading-edge protrusion velocity, under control conditions. C. Cross-correlation functions of Rac1 activity against the leading-edge protrusion velocity, under control conditions. D. Cross-correlation functions of RhoG activity against Rac1 activity, measured simultaneously, under control conditions. Cross-correlation traces are color coded as distances away from the leading-edge (see bottom of the Figure). Each trace represents an average of 27 regions and 2,916 measurement windows obtained from 12 cells across 4 independent experiments per condition. 95% confidence interval traces are not shown for clarity.

After characterizing the basic control behaviors, we next tested the effect of Src-family kinase inhibition using PP2, a broad-band Src-family kinase inhibitor (92). Src-family kinases function as key signaling components downstream of, or in close association with, activated receptor tyrosine kinases (93). They can regulate Rho-family GTPase signaling by modulating upstream regulators, including GEFs, GAPs, and RhoGDI, thereby contributing to the proper spatial and temporal coordination of Rho GTPase activities (94–96). In this context, Src-family kinase activity could promote activation of a RhoG-specific GEF, leading to RhoG activation and subsequent recruitment of the ELMO–DOCK complex, which in turn activates Rac1 in a linear signaling pathway. Alternatively, Src-family kinases may promote parallel activation of RhoG and Rac1 through distinct GEFs that target each GTPase, allowing coordinated but non-linear regulation of their activities downstream of receptor activation (13, 16, 20, 27, 45, 46, 89–91). We tested these possibilities in MEF leading-edge protrusions by inhibiting Src-family kinase activity and applying morphodynamics analysis to distinguish between these two potential models of RhoG–Rac1 coordination. Under PP2 treated conditions, the positive cross-correlation between RhoG activity and protrusion velocity was reduced relative to control cells (Fig. 5A). The correlation peak shifted slightly toward negative time lags to suggest RhoG activities are now slightly following the start of protrusive initiation, although the 95% confidence interval still spanned zero (Fig. S5C) to indicate a direct concurrence in time. At positions 6–9 and 9–12 pixels (1.8 – 2.7 and 2.7 – 3.6 µm) from the edge, the peak positions shifted further toward negative time lags relative to the maximal peak at 3–6 pixels (0.9 – 1.8 µm), and the peak cross-correlation coefficient values dipped below the significance levels (Fig. 5A). In contrast to RhoG, Src-family kinase inhibition strongly attenuated the cross-correlation functions between Rac1 activity and protrusion velocity at the leading-edge and in regions behind the edge (Fig. 5B). These results indicate that Src-family kinase inhibition alters the relationship of both RhoG and Rac1 activities to protrusive edge motion, with a stronger effect on Rac1 than on RhoG. In contrast, the autocorrelation functions of protrusion velocity were not different between control and PP2-treated cells (Fig. S5D, E), indicating that the basic fluctuation dynamics of edge protrusion were largely preserved. Thus, the differentially altered coupling of RhoG and Rac1 activities to the edge-motion under PP2 treatment is likely to reflect changes in the underlying molecular signaling landscape rather than changes in the physical dynamics of protrusion and retraction. We next directly compared RhoG and Rac1 biosensor activities in the same cells, independent of protrusive edge velocity. Under Src-family kinase inhibition, the positive RhoG – Rac1 cross-correlation observed under control conditions was largely lost, with correlation values tending toward negative values in many regions (Fig. 5C). These results suggest that Src-family kinase activity is required for the coordinated dynamics of RhoG and Rac1 at leading-edge protrusions, with a particularly strong effect on Rac1 activity dynamics.

**Figure 5:**
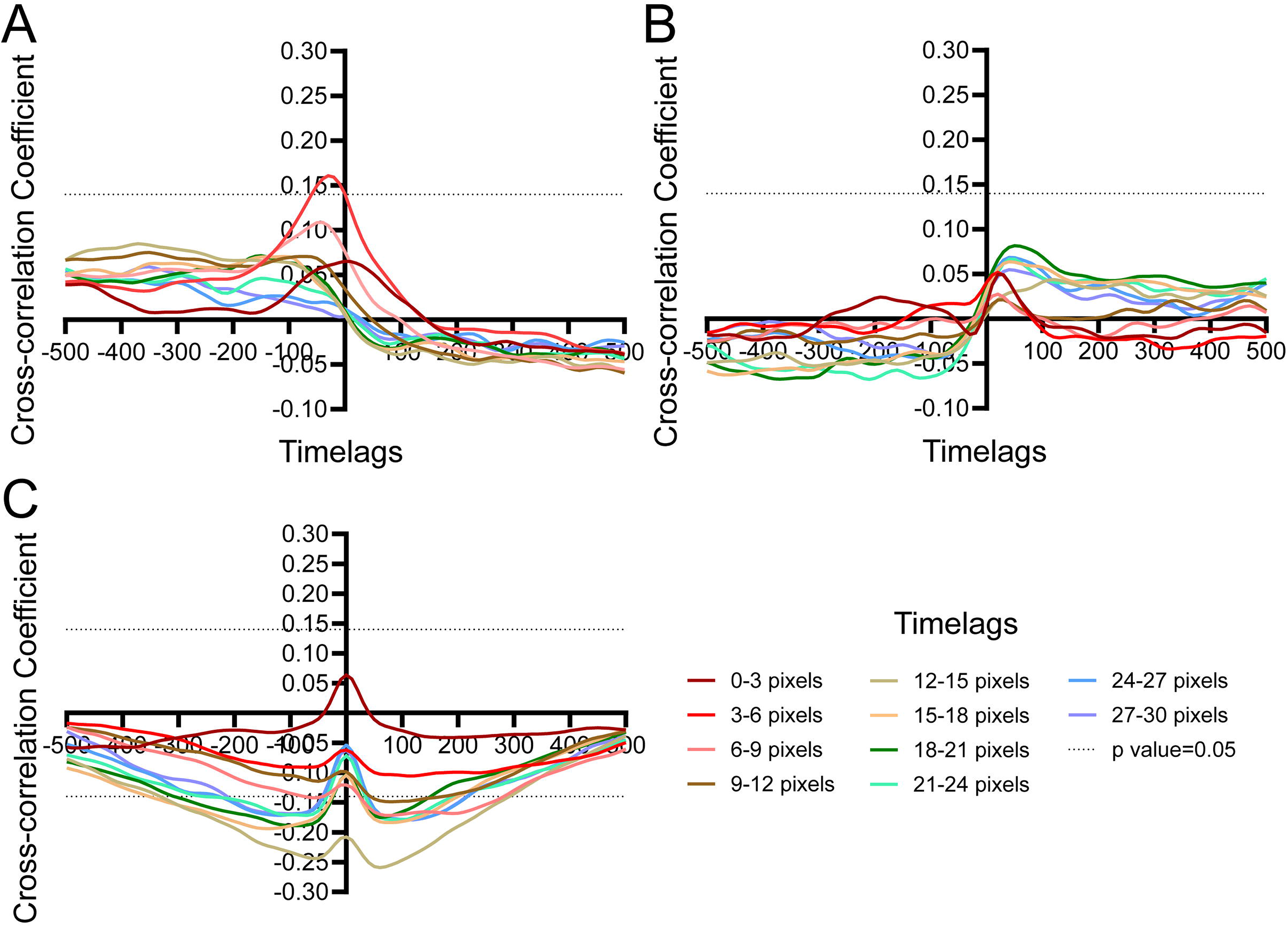
Morphodynamics mapping and cross-correlation analysis between RhoG and Rac1 activities within the leading-edge protrusions in MEF, as distances away from the edge and with Src family kinase inhibitor treatment. A. Cross-correlation functions of RhoG activity against the leading-edge protrusion velocity, under PP2 treatment (20 µM). B. Cross-correlation functions of Rac1 activity against the leading-edge protrusion velocity, under PP2 treatment (20 µM). C. Cross-correlation functions of RhoG activity against Rac1 activity, measured simultaneously, under PP2 treatment (20 µM). Cross-correlation traces are color coded as distances away from the leading-edge (see bottom of the Figure). Each trace represents an average of 27 regions and 2,916 measurement windows obtained from 12 cells across 4 independent experiments per condition. 95% confidence interval traces are not shown for clarity.

To further analyze local signaling dynamics, we applied signaling microdomain (MD) analysis (68), which identifies spatially and temporally persistent local biosensor activity clusters (Fig. 6; Supplementary Movie 5, 6). Briefly, the algorithm integrates measurements of local “coordination” of a signaling activity as determined by dynamic time warping (DTW) to create scores of coherent behavior across multiple spatial scales from which MDs can be extracted (Fig. 6A, B) (68). Unlike the standard cross-correlation type approaches, DTW is a versatile metric for the similarity of two time series even if they are not aligned or moving at differing speeds (97, 98). We computed MDs separately for RhoG and Rac1 activity in control and PP2-treated cells and analyzed MD abundance, cell area coverage, strength, and lifetimes. The average number of MDs and the distribution of MD sizes were similar across conditions for both Rac1 and RhoG, either with or without the Src-family kinase inhibitor (Fig. 6C, D). The relative coupling strengths within MDs were significantly higher for RhoG than for Rac1, both in control cells and in Src-family kinase-inhibited cells (Fig. 6E). One possible explanation for this difference is that Rac1 integrates a broader range of concurrent regulatory inputs than RhoG, which could increase the local heterogeneity of Rac1 activity and reduce the apparent coordination within individual Rac1 MDs. Interestingly, Src-family kinase inhibition significantly increased the lifetime of Rac1 MDs, whereas the lifetime of RhoG MDs was unchanged (Fig. 6F). This selective effect on Rac1 is consistent with Src-family kinases acting through multiple upstream regulators that contribute to the spatial and temporal organization of Rac1 activity, while RhoG MD turnover may be less dependent on Src-mediated regulatory inputs. Notably, the characteristic MD lifetimes, on the order of approximately 6 min, were comparable to the timescale of protrusion–retraction dynamics measured by protrusion-velocity autocorrelation (Fig. S5D, E). Thus, these signaling domains turn over on a timescale similar to that of their downstream morphological output, suggesting that MD dynamics may reflect the temporal organization of signaling underlying protrusion–retraction cycles. The whole-cell colocalization between RhoG and Rac1 MDs remained approximately 12% under both conditions (Fig. 6G), while the spatiotemporal colocalizations between RhoG and Rac1 MDs within the leading-edge 0 – 12 pixel band (0 – 3.6 µm) were significantly smaller compared to the rest of the cell and were also independent of Src-kinase inhibition (Fig. 6H). Together, these findings indicate that Src kinases directly influence the turnover dynamics of Rac1-containing signaling clusters but not RhoG. They also suggest that coupling between RhoG and Rac1 activities is stronger in regions away from the leading-edge and within the cell body. Overall, these observations also support a more prominent role for Src in Rac1 regulation than in RhoG regulation and point to a spatiotemporally discontinuous, complex signaling relationship between RhoG and Rac1.

**Figure 6:**
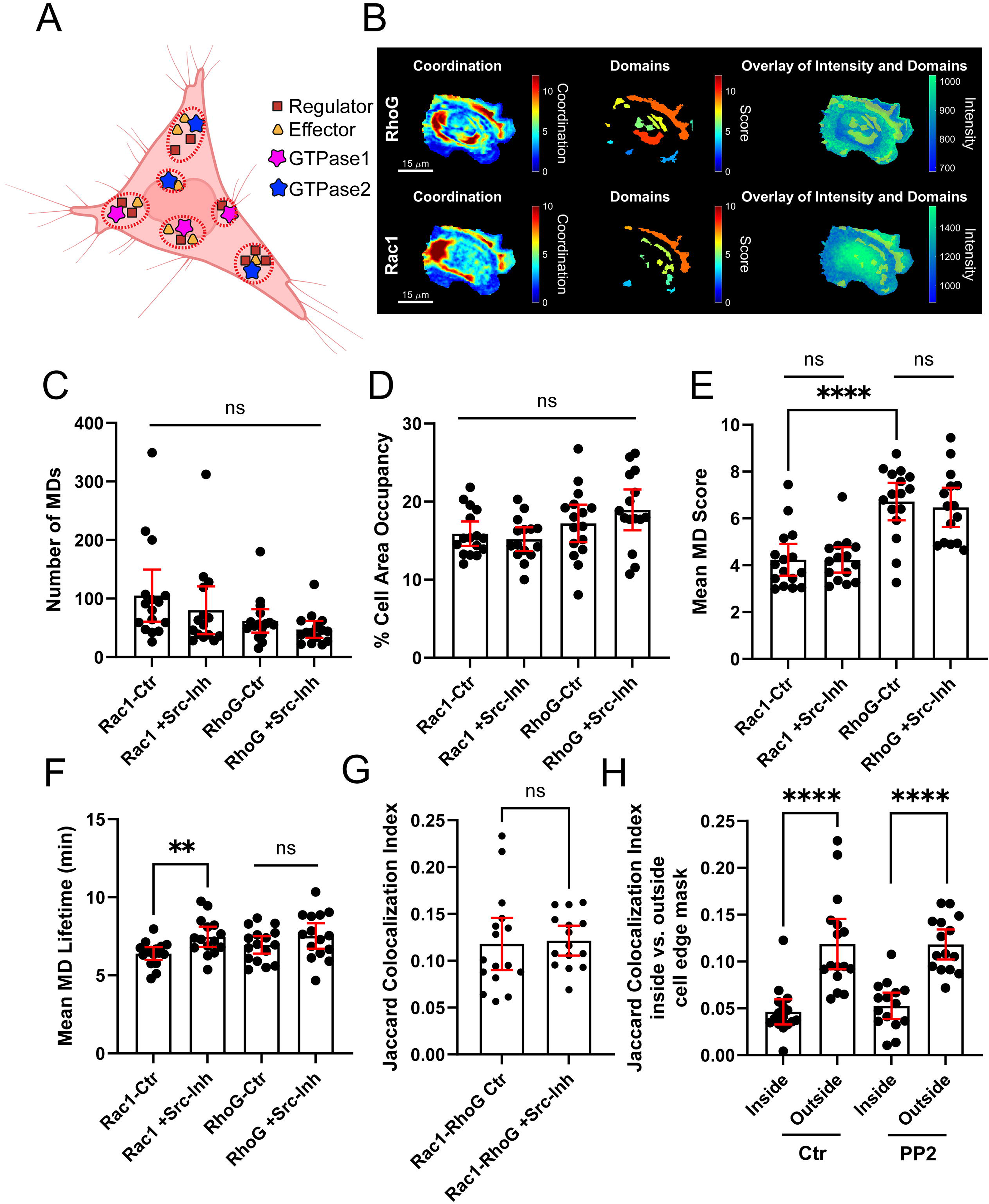
Spatial analysis of RhoG and Rac1 activities in MEFs. A. Cartoon depiction of the microdomains (MDs). A microdomain is conceptualized as a region in the cell with spatially constrained, highly coordinated Rho GTPase signaling activity that is driven by the localization of a set of GEF, GAPs, and effectors. Created with BioRender.com. B. Representative timepoint images of the computed Coordination, Domains (MD), and the overlay onto the biosensor activity Intensity map, for RhoG (TOP) and Rac1 (BOTTOM) from the control condition. C. Number of MDs per cell, n= 13-14 cells across 4 independent experiments, shown with +/- 95% confidence intervals. D. % cell area occupancy of MDs for each condition, averaged over time, n= 13-14 cells across 4 independent experiments, shown with +/- 95% confidence intervals. E. Mean MD Score, representing the strengths of the DTW coupling during MD evolution. n= 13- 14 cells across 4 independent experiments, shown with +/- 95% confidence intervals. Student’s t-test **** p<0.001. F. Mean MD lifetimes. n= 13-14 cells across 4 independent experiments, shown with +/- 95% confidence intervals. Student’s t-test ** p<0.01. G. Time averaged Jaccard colocalization index calculated from the MD analysis between Rac1 and RhoG activities, with or without the PP2 inhibitor. n= 13-14 cells across 4 independent experiments, shown with +/- 95% confidence intervals. E. Time averaged Jaccard colocalization index calculated from the MD analysis between Rac1 and RhoG activities in regions inside and outside the leading-edge 0-12 pixel region. n= 13-14 cells across 4 independent experiments, shown with +/- 95% confidence intervals. Student’s t-test **** p<0.001.

## Discussion

In this study, we developed and validated a new genetically encoded, single-chain FRET biosensor tool for RhoG and used it to define the spatiotemporal dynamics of RhoG signaling in living cells. This biosensor fills an important methodological gap in the Rho-family GTPase field. Although RhoG has been implicated in membrane trafficking, dorsal ruffling, macropinocytosis (15, 27, 99–102), and Rac1-dependent protrusive behavior (13, 16, 90), direct visualization of its activity with subcellular spatial resolution and second-scale temporal resolution has remained limited. The biosensor described here preserves full-length RhoG, including its hypervariable region and C-terminal lipid modification sites, thereby maintaining key features required for physiological membrane targeting and interaction with upstream regulators. At the same time, the use of an intramolecular ELMO1-derived affinity domain enabled robust FRET responses across multiple experimental contexts, supporting its utility as a direct readout of RhoG activation in living cells.

Several results support the specificity and regulated behavior of the biosensor. The probe showed a broad dynamic range between active and inactive RhoG conditions, responded to multiple upstream regulators, and remained sensitive to RhoGDI-mediated inhibition. The strong effect of RhoGDI3 is notable and reflects isoform-specific differences in binding, membrane extraction, or compartmental regulation of RhoG (80–82). The differential responses to panels of GAPs and GEFs further indicate that the biosensor remains accessible to relevant regulatory inputs. Importantly, although ARHGAP1 (p50RhoGAP) and ARHGAP35 (p190RhoGAP- A) have been reported to associate with an activated RhoG mutant (83, 84), no RhoG-selective GAP has yet been firmly established in a cellular context. Therefore, the effects observed in our GAP-panel analysis may reflect a combination of indirect regulation and possible direct effects on RhoG. These possibilities will require further validation using in vitro biochemical assays. Similarly, the reduction in RhoG biosensor activity readout when coexpressed with a RabGAP TBC1D10 raises the possibility of crosstalk between RhoG and membrane-trafficking pathways, either through indirect effects on membrane organization and regulator localization or through more direct regulation of RhoG activity.

A common concern for single-chain biosensors of this design is whether the affinity domain might sequester the activated GTPase or the activated GTPase within the biosensor may compete against binding endogenous downstream targets. Our data suggest that this is unlikely to be a major limitation under the conditions used here. GST pull-down experiments indicated that the activated biosensor was not readily available for promiscuous interactions with excess effector proteins when the cognate ELMO1-derived binding domain was present. This supports the interpretation that the biosensor functions primarily as a trace reporter rather than as a dominant-negative sink. Although any exogenous biosensor can perturb signaling if overexpressed, the inducible expression system and titration relative to endogenous RhoG levels helped minimize this concern (103, 104).

The accessibility of upstream GEFs and GAPs to the Rho GTPase embedded within the FRET biosensor backbone highlights an important aspect of GTPase biology that could influence biosensor readouts. Constitutively active and dominant-negative Rho GTPase mutants, including the RhoG Q61L, G12V, and T17N mutants used here, can form nonphysiological or unusually stable interactions with endogenous binding partners. The Q61L and G12V mutants are GTP-bound and incapable of GAP-stimulated hydrolysis, favoring interactions with endogenous effectors and, in some cases, regulatory proteins including GAPs (105). In contrast, dominant-negative T17N mutants preferentially interact with GEFs and could sequester them from endogenous GTPases (106, 107). These interactions are important because many endogenous GTPase-binding proteins are larger in size than the biosensor itself. Their binding could impose steric or conformational effects that alter the relative position or orientation of the donor and acceptor fluorophores. Thus, expression of mutant biosensors could produce FRET/donor ratio changes that do not solely reflect intramolecular binding between RhoG and the engineered effector domain. Our competitive pulldown experiments partially addressed this concern by showing that activated RhoG within the biosensor preferentially engaged the built-in ELMO1 RBD rather than competing for other effector binding targets. In addition, the fluorometric validation experiments were performed in a HEK293 overexpression system, in which biosensor components were expressed at levels expected to reduce the relative contribution of endogenous regulator binding to the measured FRET response. One example is the behavior of the T17N dominant-negative biosensor in the fluorometric assay (Fig. 1C). The T17N FRET/donor ratio was lower than that observed for the activated biosensor containing an incompetent binding domain, the biosensor containing effector-binding mutations, or the wild-type biosensor co-expressed with a 4-fold excess of RhoGDI3 (Fig. 1C, D). This result may reflect stable interaction of the T17N biosensor with endogenous GEFs and GEF complexes, which could reposition the FRET pair and slightly reduce FRET relative to other inactive or binding-incompetent conditions. Conversely, for the activated mutant with an incompetent intramolecular binding domain, binding by endogenous GAPs, effectors, or other protein complexes could sterically alter the biosensor conformation and modestly elevate the FRET/donor ratio. In many cells, RhoGDIs are abundant and bind Rho GTPases in a 1:1 complex, maintaining a large fraction of the inactive GTPase pool in the cytoplasm (108). Therefore, we consider the most physiologically useful dynamic range of the RhoG biosensor to be the difference between the wild-type biosensor activated by overexpression of a competent GEF fragment, such as TrioGEF (Fig. 1F), and the inactive condition produced by excess RhoGDI co-expression. For the RhoG biosensor, this dynamic range was approximately 2.5-fold. This value is likely more biologically relevant than the absolute dynamic range obtained by comparing the Q61L and T17N mutant biosensors.

Live-cell imaging revealed that RhoG activation is dynamically associated with protrusive membrane behavior, PDGF-induced dorsal ruffling, and pinocytic structure formation. These observations are consistent with prior models placing RhoG downstream of receptor tyrosine kinase signaling and upstream of actin-remodeling pathways that drive membrane protrusion and dorsal cup closure (14, 15, 27, 109). The imaging data extend these models by showing that RhoG activity is not merely present during these events, but is spatially and temporally organized within edge ruffles, dorsal ruffles, and forming pinosomes. The appearance of RhoG activity within punctate regions of developing pinocytic structures suggests that RhoG may help coordinate local signaling at the interface between actin remodeling, membrane deformation, and vesicle maturation (2, 4, 22, 47, 110).

A major advance of this study is the direct comparison of RhoG and Rac1 activities in the same living cells. RhoG has often been positioned upstream of Rac1, particularly through ELMO– DOCK complexes (13, 16, 89, 90), but can also operate in parallel (20, 27, 45, 46, 91). Our simultaneous imaging data show that both RhoG and Rac1 activities are positively coupled to protrusion, but their relationship is not consistent with a simple linear cascade. Direct RhoG– Rac1 cross-correlation was spatially heterogeneous, with strong coupling at the immediate leading edge, substantial decoupling within the frontal protrusive region, and increased coupling again farther from the edge. Thus, rather than exhibiting a simple monotonic relationship with distance from the plasma membrane, RhoG and Rac1 appear to occupy spatially distinct zones of signaling coordination. The reduced coupling within the frontal protrusive region may reflect differences in membrane compartmentalization, effector engagement, or localization of upstream regulators. RhoG may function preferentially within membrane-trafficking or membrane-remodeling contexts (99–101), whereas Rac1 is more directly integrated with actin-polymerization networks that sustain lamellipodial extension (111). The renewed RhoG–Rac1 coupling farther from the edge could reflect convergence of these pathways within more central trafficking or signaling compartments, an interpretation that is also consistent with the greater RhoG–Rac1 microdomain colocalization observed outside the leading-edge region. However, because the innermost measurement windows can approach perinuclear regions, where biosensor fluorescence distributions and differences in cellular geometry may affect ratiometric measurements, we interpret this increase in coupling cautiously. Further analysis would be valuable to determine whether the increased coupling toward the cell interior represents a distinct biological signaling compartment. Overall, the data support a model in which RhoG and Rac1 are coordinated components of a broader protrusion-associated signaling network, but their coupling varies substantially across subcellular space. Src-family kinase inhibition further refined this model. PP2 reduced the coupling of RhoG activity to protrusion but had a much stronger effect on Rac1, nearly abrogating Rac1 activity–protrusion coupling. This asymmetry suggests that Src-family kinases are not simply required for bulk activation of both GTPases. Instead, RhoG can remain partially engaged with protrusion-associated signaling in the absence of Src-family kinase activity, whereas appropriate Rac1 activity dynamics are strongly Src dependent (20, 27, 112–116). The inversion of the RhoG–Rac1 cross-correlation from positive under control conditions to negative after Src-family kinases inhibition further indicates that Src-family kinases help maintain normal coordination between these two GTPases. When Src-family kinase signaling is disrupted, RhoG and Rac1 become uncoupled or inversely organized in time, consistent with Src-family kinases acting at multiple levels of the RhoG–Rac1 network, with a particularly strong influence on Rac1 dynamics.

The signaling microdomain analysis provided further evidence for this differential Src dependence. Src inhibition did not substantially alter the number or size of RhoG or Rac1 microdomains, but it significantly prolonged Rac1 microdomain lifetimes while leaving RhoG microdomain lifetimes unchanged. This suggests that Src-family kinases do not primarily control the formation or bulk distribution of signaling clusters, but instead contribute to the turnover dynamics of Rac1-containing domains. Such selective regulation is consistent with Rac1 functioning as a highly convergent signaling node whose spatial organization is controlled by multiple upstream regulators, several of which are directly or indirectly influenced by Src-family kinases. In contrast, the unchanged RhoG MD lifetimes suggest that the mechanisms governing RhoG domain turnover are less dependent on Src-family kinase activity. Interestingly, the characteristic lifetimes of these signaling microdomains were on the order of approximately 6 min, comparable to the timescale of the protrusion–retraction dynamics measured by protrusion-velocity autocorrelation. This correspondence is not necessarily expected, as the MD analysis measures the persistence of locally coordinated molecular signaling whereas the autocorrelation analysis measures the temporal organization of a morphological output. Their similar timescales therefore suggest that the formation and turnover of localized GTPase signaling domains may be temporally matched to the protrusion–retraction cycles they regulate. Importantly, Src-family kinase inhibition prolonged Rac1 MD lifetimes without substantially changing the protrusion-velocity autocorrelation, indicating that Src-family kinase perturbation alters the molecular organization underlying protrusion without equivalently changing the characteristic timescale of the morphological cycle.

The relative coupling strengths within RhoG microdomains were also consistently higher than those within Rac1 microdomains under both control and Src-inhibited conditions. One possible explanation is that Rac1 integrates a broader range of concurrent regulatory inputs than RhoG. The convergence of multiple inputs onto Rac1 could generate greater local spatial and temporal heterogeneity in its activity, resulting in lower apparent coordination within individual Rac1 microdomains. In contrast, RhoG may operate within a more restricted set of regulatory pathways or subcellular contexts, producing more locally coherent activity domains. Although this interpretation remains speculative, it is consistent with the greater sensitivity of Rac1 dynamics to Src-family kinase inhibition observed in our morphodynamic and microdomain analyses. The relatively low colocalization between RhoG and Rac1 microdomains, especially within the frontal leading-edge band, further supports the conclusion that these GTPases share only a subset of signaling domains. Together with the cross-correlation analysis, these findings place RhoG in a spatially distinct signaling framework that connects protrusion, membrane trafficking, and Rac1 regulation without reducing RhoG to a simple upstream trigger for Rac1. Further work will be needed to identify the regulatory inputs and molecular intermediates underlying the differential organization of RhoG and Rac1 signaling.

In summary, we report a new live-cell biosensor that enables direct visualization of RhoG activity and reveals that RhoG signaling is dynamically coupled to membrane protrusion, dorsal ruffling, and pinocytic activity. Simultaneous imaging of RhoG and Rac1 shows that these GTPases are coordinated but not uniformly coupled in space and time, supporting a compartmentalized signaling relationship rather than a simple linear cascade. Finally, Src-family kinase perturbation demonstrates that Src activity is critical for maintaining normal RhoG–Rac1 coordination, with Rac1 dynamics being especially sensitive. These findings establish a new tool for studying RhoG biology and provide a framework for understanding how RhoG contributes to the spatiotemporal organization of protrusion-associated signaling networks.

## Materials and Methods

### RhoG biosensor and fluorometry

RhoG biosensor was created by linking monomeric, circularly permutated (mcp229) ECFP to a RhoG binding domain from ELMO1 (a.a. 1-115), followed by a 42 a.a. flexible linker, a monomeric circularly permutated (mcp157) Citrine-YFP, a 10 a.a. flexible linker, and a full-length wildtype RhoG. (Sequence information in Supplementary Data). Synonymous codon ver1.0 modified (117) mcp229ECFP was produced using 2-step PCR using primer set: 5’-GCTGCATATACAAGGATCCGGCATGATTACCCTGGGAATGGATGAACTGTATAAA-3’, 5’-TTGGACACCATGCCGCCGCTGCCGCCTTTATACAGTTCATCCATTCCCA-3’, and 5’-ACTGTATAAAGGCGGCAGCGGCGGCATGGTGTCCAAAGGAGAAGAACTGTT-3’, 5’-GGTTAATACATGTTAGAAGCTTCCGGCAGCTGTCACAAATTCCAGC-3’, and then amplified using the primer pair: 5’-GGAATTATAATTATACCATGGGCATTACCCTGGGAATGGATGAACT-3’ and 5’-CGTATATTAAAATTTGGATCCTCCGGCAGCTGTCACAAATTCCAGCA-3’ and ligated into NcoI/BamHI sites of the biosensor backbone. The linker segment consisted of 2 units of repeating, flexible linker of optimized design produced and used previously (69), which was ligated into the biosensor backbone at HindIII/NotI restriction sites. Mcp157 Citrine-YFP amplified using 2-step PCR, with the primer set: 5’-GGTTATACATTATAAGCGGCCGCTATGCAGAAGAACGGCATCAAGGTGA-3’, 5’-GCCACCGCTGCCACCCTTGTACAGCTCGTCCATGCCGAGA-3’, and 5’-GCTGTACAAGGGTGGCAGCGGTGGCATGGTGAGCAAGGGCGAGGAGCTGT-3’, 5’-CCTATTATTAATAATGAATTCCTTGTCGGCCATGATATAGACGTT-3’ and then the terminal linker region was extended using the 2-step PCR reverse primer set 5’-GGTTATACATTATAAGCGGCCGCTATGCAGAAGAACGGCATCAAGGTGA-3’ and 5’-CTCCGCTAGACCCGCTGCCGCTTCCCTTGTCGGCCATGATATAGACGTTG-3’ and ligated into NotI/EcoRI sites of the biosensor backbone. ELMO1 binding domain was PCR amplified using the primer pair: 5’-GGTTTAAGATTAACAAGGATCCATGCCGCCGCCTTCGGACATCGTGAA-3’ and 5’-CGTAATATATTAATTCAAGCTTCCGGTAACATCCCGGGAGAGGCTGGCCAA-3’ and ligated into BamHI/HindIII sites of the biosensor backbone. Full length human RhoG was PCR amplified using the primer pair: 5’-GGATAACTTATATTAGAATTCATGCAGAGCATCAAGTGCGTGGTGGT-3’ and 5’-CGTTATCATTAACTTAAACTCGAGTCACAAGAGGATGCAGGACCGCCCA-3’ and ligated into EcoRI/XhoI sites of the biosensor backbone. The biosensor cDNA cassette was subcloned into pTriEX-HisMyc4 (Sigma-Aldrich, St. Louis MO, USA) for transient expression. Characterization of biosensor response was performed in HEK293 cells overexpressing the various biosensor mutants and biosensor with or without the appropriate upstream regulators. Briefly, HEK293T cells were plated overnight at 9 × 10^5^ cells/well in six-well plates coated with 0.001% poly-L-lysine (Sigma-Aldrich, St. Louis MO, USA) and transfected the following day using Polyethylenimine (PEI: Poly[imino(1,2-ethanediyl)] hydrochloride; Sigma-Aldrich, St. Louis MO, USA) transfection reagent according to the published optimized procedures (118). After 48 h transfection, cells were washed once with PBS, briefly trypsinized, and resuspended in 500 µL of cold PBS per well. Cell suspensions were stored on ice until assay. Fluorescence emission spectra were measured with a spectrofluorometer (Horiba-Jobin-Yvon Fluorolog-3MF2; HORIBA, New Brunswick, NJ, USA). The fluorescence emission spectra were obtained by exciting the cell suspension in a 500 µL quartz cuvette (Starna Cells, Atascadero, CA, USA) at 433 nm, and emission fluorescence was scanned between 450–600 nm. The background fluorescence reading of cells containing an empty vector (pCDNA3.1; Thermo Fisher Scientific, Waltham, MA, USA) was used to measure light scatter and autofluorescence and was subtracted from the data. The resulting spectra were normalized to the peak of the donor mCerulean emission intensity at 474 nm to generate the final, normalized spectra. The biosensor and the regulator cDNAs were co-transfected at ratios of 1:4 for the biosensor and the GDI, 1:2 for the GAP and 1:1 for the GEF.

### Cell culture

HEK293T cell derivative LinXE cell line were cultured in Dulbecco’s modified Eagle’s medium (DMEM; Mediatech, Manassas, VA, USA) with 10% FetalGro-EX (Rocky Mountain Biologicals, Missoula, MO, USA), 100 U/mL penicillin and 100 μg/mL streptomycin (Thermo Fisher Scientific, Waltham, MA, USA). and maintained at 37°C and 5% CO_2_ atmosphere. MEF/3T3 tet-OFF (Takara, Mountain View, CA, USA) were cultured in DMEM with 10% FBS, 100 U/mL penicillin and 100 μg/mL streptomycin and maintained at 37°C and 5% CO_2_ atmosphere. A stable cell line expressing inducible RhoG biosensor was produced using the pRetro-X-Puro tet-inducible retroviral system (Takara, Mountain View, CA, USA) as previously described (77, 103). This was used also to transduce the RhoG biosensor into MEF cells stably harboring NIR Rac1 FRET biosensor (67), as previously described. To repress the biosensor expression during normal culture, 1 µg/ml doxycycline was applied. To induce RhoG biosensor expression, doxycycline was removed 48 hours prior to imaging by detaching cells through brief trypsinization and then replating at 1×10^5^ cells per 10cm dish. Cells were plated on fibronectin (Sigma-Aldrich, St. Louis MO, USA) (FN; 10 μg/ml)-coated glass coverslips at 5×10^4^ cells for 2 hours prior to imaging. Imaging was performed in FluoroBrite DMEM without phenol red (Thermo Fisher Scientific, Waltham, MA, USA) with 3% FBS in a heated closed chamber.

### Biosensor GST-pulldown

Competitive effector binding and pull-down experiments were performed as following. GST-fusion of the ELMO1 RBD (1-115 a.a.) was produced by PCR amplification using the primer pair 5’-GGTTTAAGATTAACAAGGATCCATGCCGCCGCCTTCGGACATCGTGAA-3’ and 5’-GGTTTAAATTTCAATTGGATCCGGTAACATCCCGGGAGAGGCTGGCCAA-3’, and ligated into pGEX-4T3 (GE Lifesciences, Chicago, IL, USA) at BamHI site and used for bacterial protein expression and purified using standard protocols (119). LinXE cells were plated overnight at 9 ×10^5^ cells per well of a 6 well plate and transfected using the PEI reagent at 8 µL with 2 µg/well of pCDNA3.1 (Thermo Fisher Scientific, Waltham, MA, USA) empty vector, pTriEX-mGreenLantern-RhoG Q61L, pTriEX-mGreenLantern-RhoG T17N, constitutively activated RhoG Q61L biosensor, or constitutively activated RhoG Q61L biosensor but with the ELMO1 RBD replaced with an inert binding domain from the p21 activated kinase1 binding domain (PBD) containing three additional non-binding mutations H83/86D and W103D (60, 120, 121). 24 hrs following transfection, cells were lysed in RBD-pulldown buffer (50 mM Tris pH7.4, 500mM NaCl, 50mM MgCl_2_, and 1% NP-40) containing protease inhibitor cocktail (Sigma-Aldrich, St. Louis MO, USA) and Pefabloc SC (Sigma-Aldrich, St. Louis MO, USA). The GST-RBD pulldown experiments were performed as previously described (122).

### Exogenous stimulation with PDGF

For exogenous stimulation, MEF cells induced to express the RhoG biosensor were plated on FN-coated coverslips at 5×10^4^ cells and serum starved in DMEM containing 5% BSA for 24hrs prior to stimulation with 100 ng/mL PDGF (R&D Systems, Minneapolis, MN, USA) for the indicated times at 37°C before fixation in 1.0 % formaldehyde in PBS for 15 min.

### Microscopy imaging

Activations of RhoG biosensor were measured by observing the ratio of FRET emission to the donor ECFP emission. For live cell experiments using MEF cells, images were acquired at 40X magnification (Olympus UIS2 40X N/A 1.3; Evident Scientific, Waltham, MA, USA) using a custom microscope based on Olympus IX83 ZDC microscope system (Evident Scientific, Waltham, MA, USA), capable of simultaneous acquisitions of FRET with donor emission channels, through two PrimeBSI-Express cooled sCMOS cameras (Teledyne Photometrics, Tucson, AZ, USA) mounted via a beamsplitter (123). For the dual biosensor multiplex analysis of RhoG and Rac1, the beamsplitter system was extended to include two additional PrimeBSI cooled sCMOS cameras (Teledyne Photometrics, Tucson, AZ, USA). Cells were illuminated using a solid-state LED white-light illumination (Lumencor, Beaverton, OR, USA), and a filterwheel controlled the appropriate excitation wavelengths for the biosensors: ET436/20X (Chroma Technology, Bellows Falls, VT, USA) for ECFP and FF01-628/32 (AVR-Optics, Fairport, NY, USA) for miRFP670. For the emission paths, following filters were used: ET480/40M (Chroma Technology, Bellows Falls, VT, USA) for ECFP; ET535/30M (Chroma Technology, Bellows Falls, VT, USA) for FRET Citrine-YFP; FF01-681/24 (AVR-Optics, Fairport, NY, USA) for miRFP670; and FF01-794/160 (AVR-Optics, Fairport, NY, USA) for miRFP720. For dichroic mirrors, the main dichroic mirror turret in the microscope contained ZT440/488/561/635rpc-UF1 (Chroma Technology, Bellows Falls, VT, USA). In the emission port of the beamsplitter, T560LPXRXT-UF2 (Chroma Technology, Bellows Falls, VT, USA) separated the cyan-yellow wavelengths from red and NIR wavelengths, followed by T505LPXR-UF2 (Chroma Technology, Bellows Falls, VT, USA) which splitted the donor ECFP from FRET Citrine-YFP emissions. For the splitting of miRFP670nano1 from miRFP720 wavelengths, T700LPXR-UF2 (Chroma Technology, Bellows Falls, VT, USA) was used. Image acquisitions at 4 cameras and the microscope motion control functions were all controlled by Visiview version 7.0.0.9 (Visitron Systems, Puchheim, Germany). The simultaneous camera operations were synchronized using an external Virtex trigger system (Visitron Systems, Puchheim, Germany) controlled by Visiview. Each FRET and donor Image pairs were simultaneously acquired and were properly aligned for a pixel-by-pixel matching using a priori calibration and affine coordinate transformation-based morphing to achieve accurate registration prior to ratiometric calculations as described previously (124). Image processing, including flat-field correction, background subtraction, ratio calculations, and the correction for photobleaching were performed as described previously (124). For fixed cell imaging, a 60X magnification objective lens (Olympus UIS2 60X N/A 1.5; Evident Scientific, Waltham, MA, USA) was used. In both live-cell imaging and fixed-cell imaging, DIC transilluminated images were also obtained.

For fluorescence lifetime imaging microscopy (FLIM), we used the FastFLIM system (Inscoper, Cesson-Sévigné, France). Briefly, this system enables rapid determination of relative, pixel-wise mean fluorescence lifetimes without requiring exponential fitting or calibration against a sample with known fluorescence lifetime characteristics (125–129). Unlike TCSPC or frequency-domain FLIM approaches, FastFLIM does not directly measure absolute fluorescence lifetimes. Instead, it estimates the mean fluorescence lifetime at each pixel from a series of time-gated intensity images. This is achieved by acquiring narrow excitation and emission gates, typically 100–2000 ps wide, within a 10 ns time window. Five to ten consecutive gated images are then used to calculate the mean fluorescence lifetime at each pixel according to the following equation: τ = ΣΔt\ × I\ / ΣI\, where Δt\ is the delay time of the ith gate and I\ is the corresponding pixel-wise time-gated intensity image (125, 126, 128, 129). FLIM was performed on a custom Olympus IX83 microscope system (Evident Scientific, Waltham, MA, USA) using a 100×, 1.5 NA objective lens. Imaging conditions were selected such that the gate widths for successive acquisitions were 500–1000 ps, providing greater than 2.5-fold temporal sampling relative to the expected fluorescence lifetime of the donor fluorophore. For each field of view, 10 sequential gated frames were acquired at these gate widths to determine the relative, pixel-wise mean fluorescence lifetime. In addition, five consecutive lifetime image sets were acquired and averaged for each field of view to reduce random measurement noise. The donor-only condition, mcp229ECFP, was used as the baseline for calculating lifetime changes, reported as delta lifetime values.

### Edge tracking and cross-correlation analysis

The Morphodynamics algorithm was used to track and construct the measurement windows as previously described (39). Briefly, the leading-edge tracking was performed on a segment of a cell image movie that showed robust protrusion/retraction cycling. The cropped image stack was processed to track the edge motion using the prPanel.m protrusion tracking software (39). The measurement windows of the size 3 by 6 pixels which translated to 0.927μm by 1.854μm at 40x magnification was taken previously to be the diffusion-limited area size for the 10s time intervals of each successive time points of acquisition. Typically, the entire leading-edge segment measured contained 30 – 100 measurement windows depending on the overall length of the segment. The window positions were successively moved back and away from the leading-edge in 3-pixel units, to calculate the spatial dependence of the cross-correlation functions. This was used to measure the correlational coupling up to 10 μm distance from the leading-edge. The normalized cross-correlation coefficient was computed at each window between the measured velocity in the normal direction at the edge and the changes in RhoG or Rac1 activities at the corresponding window, or in case of direct multiplex measurements, of RhoG and Rac1 activities, using the Matlab function xcov. The individual cross correlation coefficient distribution at each window was treated as an independent measurement entity, smooth-spline fitted, pooled between all cells imaged and the average maximal cross-correlation coefficient timelag location and the 95% confidence interval were calculated by a non-parametric bootstrap method (130). For velocity auto-correlation functions, the same procedure was used but the protrusion edge velocities were cross-correlated against itself.

### Signaling microdomains analysis

Signaling microdomains (MDs) were detected and followed over time using the previously described microdomain analysis framework (52). For each intensity time-lapse sequence, a matching coordination movie was generated by combining the local temporal coordination measured at each pixel with that of neighboring pixels across a defined range of downsampling scales and bandwidth parameters. This coordination movie serves as a dynamic map of local signal coordination, in which spatially restricted regions of high coordination correspond to signaling microdomains, as illustrated in Fig. 5A. Candidate MDs were obtained by segmenting the coordination movie, after which weakly scored regions were discarded. The remaining candidate regions were then linked across successive frames to define the final set of spatiotemporal microdomains. A full description of this procedure is provided in (52).

MD analyses were carried out using characteristic domain lifetimes of 10, 25, 51,76, 101, 151, and 175 frames, with image acquisition performed every 10 s. A minimum area threshold of 40 connected pixels was required, whereas no maximum size threshold was used. Pixel connectivity in the x-y plane was defined by 4-neighbor adjacency. Overlap between MDs was quantified with the Jaccard index, J(A, B) = (A \ B)/(A ∪ B) (131).

## Supplementary Figure Legends

Figure S1: Characterization of RhoG biosensor. A. Representative Western blots showing GST-RBD pull-downs of RhoG mutants, anti-GFP detections are shown. The GST fusion construct of RhoGTPase binding domain from ELMO1 (a.a. 1-115) was used to pull-down on mGreenLantern fusion RhoG mutants. Lanes: 1 untransfected control; 2 wildtype RhoG; 3 Q61L RhoG; 4 G12V RhoG; 5 F28L RhoG; and 6 T17N RhoG. B. Same set of blots as in A, showing the loading controls. C. Spectrofluorometric analysis of the RhoG biosensor in suspended cells, in response to increasing concentrations of the co-transfected GDI1. Data represent mean ± SEM. Statistical significance was determined using unpaired two-tailed Student’s t-tests. *p<0.05, **p<0.01, from n=4-13 independent experiments. D. Spectrofluorometric analysis of the RhoG biosensor in suspended cells, in response to increasing concentrations of the co-transfected GDI2. Data represent mean ± SEM. Statistical significance was determined using unpaired two-tailed Student’s t-tests. *p<0.05, **p<0.01, from n=4-13 independent experiments. E. Spectrofluorometric analysis of the RhoG biosensor in suspended cells, in response to increasing concentrations of the co-transfected GDI3. Data represent mean ± SEM. Statistical significance was determined using unpaired two-tailed Student’s t-tests. *p<0.05, **p<0.01, from n=4-13 independent experiments.

Figure S2: Representative, time-lapse panel of RhoG activity in a MEF cell, in response to PDGF stimulation and undergoing macropinocytosis. Whole cell views: Scale bar = 10 µm. Zoomed views: Scale bar = 2 µm. Example macropinosome is shown with black arrows. Warmer pseudocolor scale corresponds to higher RhoG activity.

Figure S3: Biosensor expression levels in stably transduced MEF cells. A. Western blot of RhoG biosensor expression in MEFs under tet-OFF induced expression upon removal of Doxycycline. B. Western blot of NIR Rac1 biosensor expression in MEFs under tet-OFF induced expression upon removal of Doxycycline. C. β-actin loading control for A and B. D. Quantification of relative biosensor expression levels normalized against the corresponding endogenous GTPase expression levels. Data are shown as means +/-95% confidence intervals, of N=3 independent experiments. E. Quantification of endogenous RhoG and Rac1 expression levels with or without the biosensor induction. Data are shown as means +/-95% confidence intervals, of N=3 independent experiments.

Figure S4: Morphodynamics mapping and cross-correlation analysis between RhoG and Rac1 activities within the leading-edge protrusions in MEF, as distances away from the edge and with or without Src family kinase inhibitor treatment, shown as the peak average cross-correlation coefficient distribution at each distance segments. A. Peak cross-correlation coefficient values between the leading-edge protrusion velocity and RhoG activity, as a function of the distance away from the leading-edge. Both control (LEFT) and the PP2 treated (20 µM) (RIGHT) conditions are shown. B. Peak cross-correlation coefficient values between the leading-edge protrusion velocity and Rac1 activity, as a function of the distance away from the leading-edge. Both control (LEFT) and the PP2 treated (20 µM) (RIGHT) conditions are shown. C. Peak cross-correlation coefficient values between RhoG activity and Rac1 activity, measured simultaneously, as functions of distances away from the leading-edge. Both control (LEFT) and the PP2 treated (20 µM) (RIGHT) conditions are shown. Cross-correlation coefficient bars are color coded as distances away from the leading-edge (mirroring data shown in Figure 4). Data are shown as means +/- 95% confidence intervals. Data represents an average of 27 regions and 2,916 measurement windows obtained from 12 cells across 4 independent experiments per condition. Student-t test: * p<0.05, *** p<0.001, **** p<0.0001.

Figure S5: A. Zoomed in view showing the cross-correlation function traces of the leading-edge velocity against the RhoG activity from the first two segments, shown with +/- 95% confidence intervals. Control condition is shown, corresponding to the plot shown in Figure 4A. B. Zoomed in view showing the cross-correlation function traces of the leading-edge velocity against the RhoG activity from the first two segments, shown with +/- 95% confidence intervals. PP2 (20 µM) treated condition is shown, corresponding to the plot shown in Figure 4B. B. Zoomed in view showing the cross-correlation function traces of the leading-edge velocity against the Rac1 activity from the first two segments, shown with +/- 95% confidence intervals. Control condition is shown, corresponding to the plot shown in Figure 4C. D. Average auto-correlation function of the leading-edge protrusion velocities under control conditions, shown with +/- 95% confidence intervals. E. Average auto-correlation function of the leading-edge protrusion velocities under PP2 (20 µM) treated conditions, shown with +/- 95% confidence intervals.

## Description of Additional Supplementary Files

Supplementary Movie 1. Control MEF cell expressing RhoG biosensor, showing DIC (LEFT) and RhoG activity (RIGHT). Cell was imaged using a 40× objective and 10\s acquisition intervals. Movie is displayed at a playback rate of 7 frames per second (fps). Scale bar, 20\μm.

Supplementary Movie 2. Zoomed in view of the leading-edge protrusion from a control MEF cell expressing RhoG biosensor, showing DIC (LEFT) and RhoG activity (RIGHT). Cell was imaged using a 40× objective and 10\s acquisition intervals. Movie is displayed at a playback rate of 7 frames per second (fps). Scale bar, 5\μm.

Supplementary Movie 3. Zoomed in view of the edge ruffling resulting in pinocytosis from a PDGF (100 ng/mL) stimulated MEF cell expressing RhoG biosensor, showing the FRET donor channel (LEFT) and RhoG activity (RIGHT). Cell was imaged using a 40× objective and 10\s acquisition intervals. Movie is displayed at a playback rate of 7 frames per second (fps). Scale bar, 5\μm.

Supplementary Movie 4. Zoomed in view of the macropinocytosis from a PDGF (100 ng/mL) stimulated MEF cell expressing RhoG biosensor, showing the RhoG activity (LEFT) and DIC (RIGHT). Cell was imaged using a 40× objective and 10\s acquisition intervals. Movie is displayed at a playback rate of 7 frames per second (fps). Scale bar, 2\μm.

Supplementary Movie 5. Representative RhoG and Rac1 signaling MD analysis under control conditions. Representative time-series images of the biosensor activities, signaling coordination strength, MDs, and biosensor activity-MD overlay are shown, for RhoG (TOP) and Rac1 (BOTTOM). Images were taken at 10 s intervals. Frame rate 7 fps. White bar = 15 µm.

Supplementary Movie 6. Representative RhoG and Rac1 signaling MD analysis under PP2 (20 µM) treated conditions. Representative time-series images of the biosensor activities, signaling coordination strength, MDs, and biosensor activity-MD overlay are shown, for RhoG (TOP) and Rac1 (BOTTOM). Images were taken at 10 s intervals. Frame rate 7 fps. White bar = 15 µm.

Supplementary Data: Base-pair sequence of Cyan-Yellow RhoG biosensor.

Supplementary Text: Engineering and optimization details of the single-chain RhoG FRET biosensor.

## Author Contributions

M.d.A.L, A.T., V.M., and L.H. performed the experiments. R.R. wrote and optimized the computational pipelines. R.G.M. evaluated the binding domain efficacy. G.D. supervised the computational analysis. R.G.M. and L.H. conceived the project. R.G.M., G.D., V.M., D.C., and L.H. reviewed the results and analysis. M.d.A.L. and L.H. wrote and revised the manuscript. All authors have read and agreed to the published version of the manuscript.

Institutional Review Board Statement: Not applicable for studies not involving humans or animals. Document of Registration #250032 for Recombinant DNA, Albert Einstein College of Medicine, Department of Environmental Health and Safety.

## Informed Consent Statement

Not applicable for studies not involving humans.

## Data Availability Statement

The raw data supporting the conclusions of this article will be made available by the authors on request. The biosensor expression constructs are available from the authors on request. The image processing tools and computational algorithms are available from the authors on request.

## Acknowledgments

This work was supported by NIH grants R35GM136226 (L.H.), R01GM136826 and R15GM155874 (R.G.M), R35GM162265 (V.M.), and the Chan Zuckerberg Initiative MET-0000000459 (L.H.). L.H. is an Irma T. Hirschl Career Scientist. Support by the Danuser lab for this work was funded by RM1GM145399. FACS sorting was conducted in the Albert Einstein College of Medicine Flow Cytometry Facility, funded in part by the NCI Comprehensive Cancer Center grant P30CA013330. We thank members of the Segall and Cox laboratories at Albert Einstein College of Medicine for their helpful discussions.

