## SupplementaryText for "A genetically encoded RhoG FRET biosensor reveals spatially compartmentalized RhoG–Rac1 signaling during cell protrusion"

**Supplementary Text: Engineering of the RhoG biosensor**

**Biosensors based on the mCerulean–mVenus FRET pair**

The first single-chain RhoG biosensor was designed using previously established FRET biosensor platforms. The first construct was modeled after the ECFP–Citrine RhoA biosensor (38). We designated this configuration "Type I," based on the position of the RhoGTPase-binding domain relative to the first fluorescent protein (FP1). From the N- to the C-terminus, the biosensor consisted of the ELMO1 RhoG-binding domain (BD; amino acids 1–115) (4), mCerulean (mCer) (132), a flexible linker of optimized length (133), mVenus (mVn) (134), and full-length human RhoG (Fig. ST1A). The linker consisted of one to four repeating 17-amino-acid units, designated Linkers 1–4.

We first compared the constitutively active G12V and WT versions of the biosensor in live HEK293 cell suspensions using spectrofluorometry. To determine whether RhoG inhibition altered the biosensor

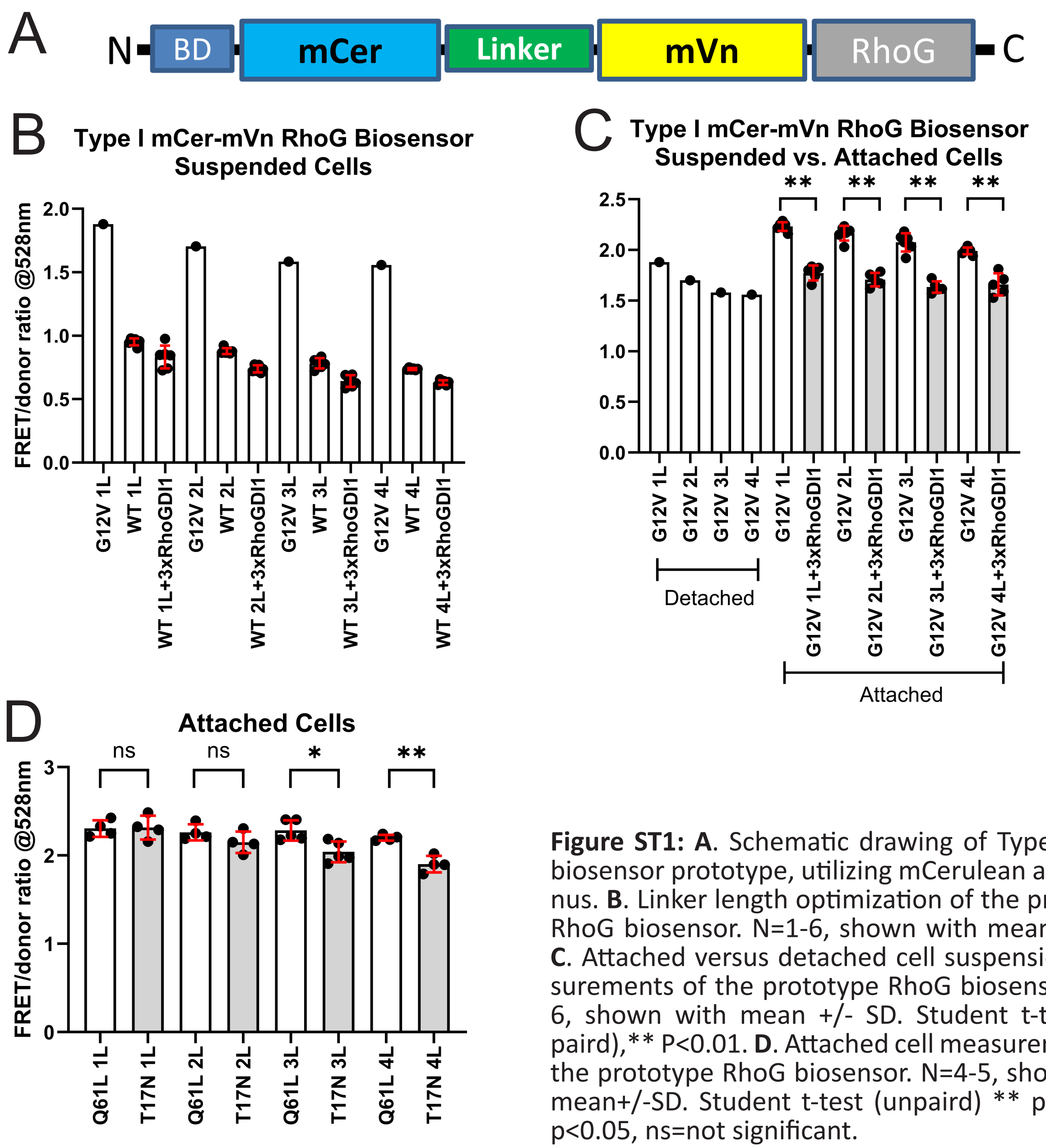


**Figure ST1: A**. Schematic drawing of Type I RhoG biosensor prototype, utilizing mCerulean and mVenus. **B**. Linker length optimization of the prototype RhoG biosensor. N=1-6, shown with mean +/- SD. **C**. Attached versus detached cell suspension measurements of the prototype RhoG biosensor. N=1-6, shown with mean +/- SD. Student t-test (unpaired),** P<0.01. **D**. Attached cell measurements of the prototype RhoG biosensor. N=4-5, shown with mean+/-SD. Student t-test (unpaired) ** p<0.01, * p<0.05, ns=not significant.

response, we also co-expressed the WT biosensor with a threefold excess of RhoGDI1. RhoGDI3 was not included in these initial experiments because a corresponding construct had not yet been generated in the laboratory. As expected, the WT biosensor exhibited a moderately lower FRET/donor ratio than the G12V mutant. Co-expression of excess RhoGDI1 further reduced the FRET/donor ratio of the WT biosensor by approximately 15%, with little apparent dependence on linker length (Fig. ST1B). By comparison, the final RhoG biosensor described in this study showed an approximately 23% difference under similar measurement conditions when the WT biosensor was co-expressed with a three- to fourfold excess of RhoGDI1 (Fig. S1C).

We next examined whether cell attachment affected the response of the G12V biosensor. Biosensors expressed in adherent HEK293 cells exhibited slightly higher FRET/donor ratios than those measured in suspended cells (Fig. ST1C). Under adherent conditions, co-expression of a threefold excess of RhoGDI1 reduced the FRET/donor ratio of the G12V biosensor by approximately 18–20%. In addition to G12V, we tested the constitutively active Q61L and dominant-negative T17N versions of the same Type I biosensor under adherent conditions. Although the Q61L construct consistently produced an elevated FRET/donor ratio, the T17N construct was not clearly distinguishable from the Q61L construct (Fig. ST1D). Because this design did not provide sufficient separation between constitutively active and dominant-negative RhoG states, further development of this Type I biosensor was discontinued.

Based on previous biosensor designs (38, 63, 64, 81, 135, 136), the limited difference in FRET/donor ratio between dominant-negative and constitutively active states could arise from at least two factors. First, the affinity domain may not adequately distinguish between high- and low-affinity states of the RhoG effector-binding interface. Alternatively, the structural orientation of the affinity domain relative to RhoG may be unfavorable, preventing nucleotide-state-dependent changes from producing an appropriate change in FRET. The Q61L and T17N response profiles supported the latter possibility (Fig. ST1D). Increasing the length of the intramolecular linker tended to improve separation between the active and inactive biosensor states. The additional linker length may provide greater conformational freedom, allowing the two FRET moieties to adopt orientations that more effectively distinguish between nucleotide states. These findings suggested that suboptimal structural alignment, rather than an inherent inability of the affinity domain to discriminate between RhoG activation states, was likely the main cause of the limited biosensor response.

Based on this analysis, we next sought to optimize both the RhoG-binding domain and the relative orientation of the biosensor components. A longer ELMO1 BD fragment comprising amino acids 1–315 was previously shown to support selective pulldown of RhoG (137). We therefore extended the ELMO1 BD in our biosensor to amino acids 1–330. In parallel, we generated a second biosensor configuration, designated “Type II”, based on our cyan–yellow Rac1 FRET biosensor design (64). In this orientation, the positions of FP1 and the BD were switched relative to the Type I design (Fig. ST2A).

We tested the constitutively active Q61L and dominant-negative T17N versions of these biosensors, each containing one to five linker units, in detached and suspended HEK293 cells (Fig. ST2B, C). The Type I orientation produced little or no significant separation between the two mutant conditions. In some

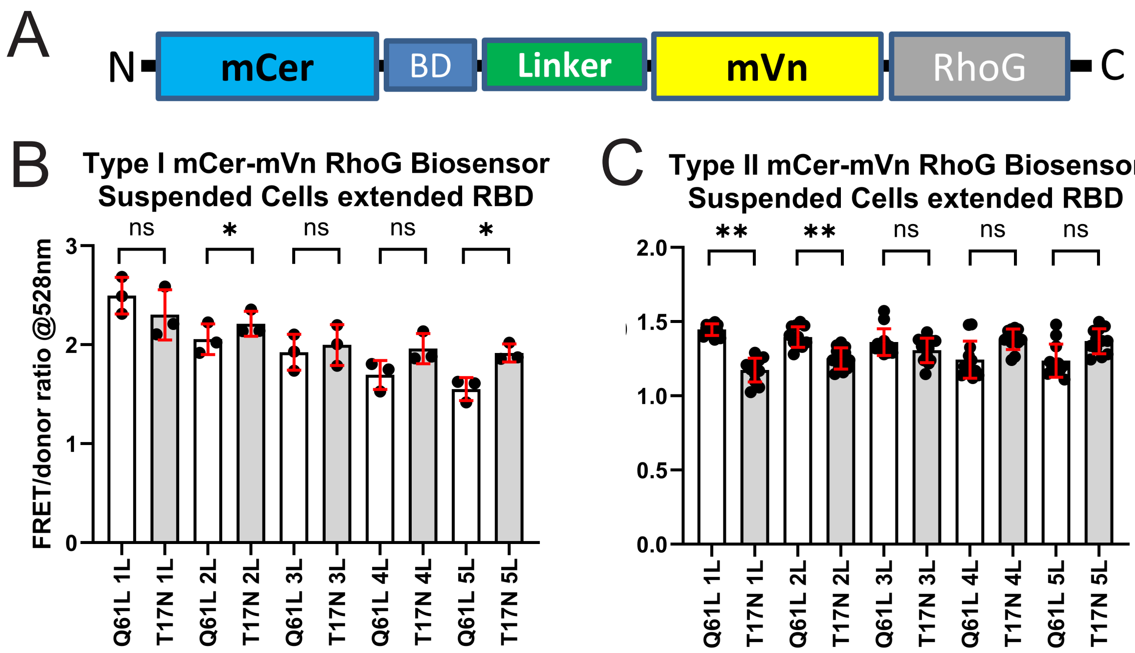


**Figure ST2: A**. Schematic drawing of Type II RhoG biosensor prototype, utilizing mCerulean and mVenus. **B**. Type I biosensor as in Figure ST1A but with an extended ELMO1 RBD (1-330 amino acids). Detached cell suspension fluorormetric measurements with various linker lengths. N = 3, shown with mean +/- SD. Student-te test (unpaired), * $p<0.05$, ns = not significant. **C**. Type II biosensor with an extended ELMO1 RBD (1-330 amino acids). Detached cell suspension fluorormetric measurements with various linker lengths. N = 12, shown with mean +/- SD. Student t-test (unpaird) ** $p<0.01$, ns=not significant.

linker variants, the response was slightly inverted, with the T17N mutant showing a modestly higher FRET/donor ratio than the Q61L mutant. By contrast, although the absolute FRET/donor ratios of the Type II biosensors were generally lower than those of the Type I biosensors, several Type II linker variants produced stronger separation between the Q61L and T17N states. Importantly, in the Type II orientation, the T17N mutants showed reduced FRET/donor ratios, consistent with the expected direction of biosensor deactivation (except in longer linkers, i.e., 4 and 5 linkers). These results indicated that the Type II configuration provided a more favorable orientation for detecting RhoG nucleotide-state-dependent changes.

We next asked whether the difference in FRET/donor ratio between the Q61L and T17N versions of the Type II biosensor could be further improved by altering the dipole-coupling geometry between the FRET donor and acceptor fluorescent proteins. To do this, we modified the acceptor fluorescent protein by circular permutation. We tested previously described circularly permuted mVenus variants, including cp49, cp157, cp173, cp195, and cp229 mVenus (74). These variants were introduced into the one-linker Type II biosensor, which had produced the largest difference between the Q61L and T17N responses in the previous analysis. The resulting biosensors (Fig. ST3A) were tested for changes in FRET/donor ratio in detached, suspended HEK293 cells. Compared with the original non-permuted mVenus acceptor, none of the circularly permuted acceptor variants improved separation between the active and inactive biosensor states. The largest difference remained approximately 41%, which was observed with the original fluorescent protein FRET pair (Fig. ST3B).

Based on the results shown in Fig. ST2C, the Type II biosensor containing the extended ELMO1 BD

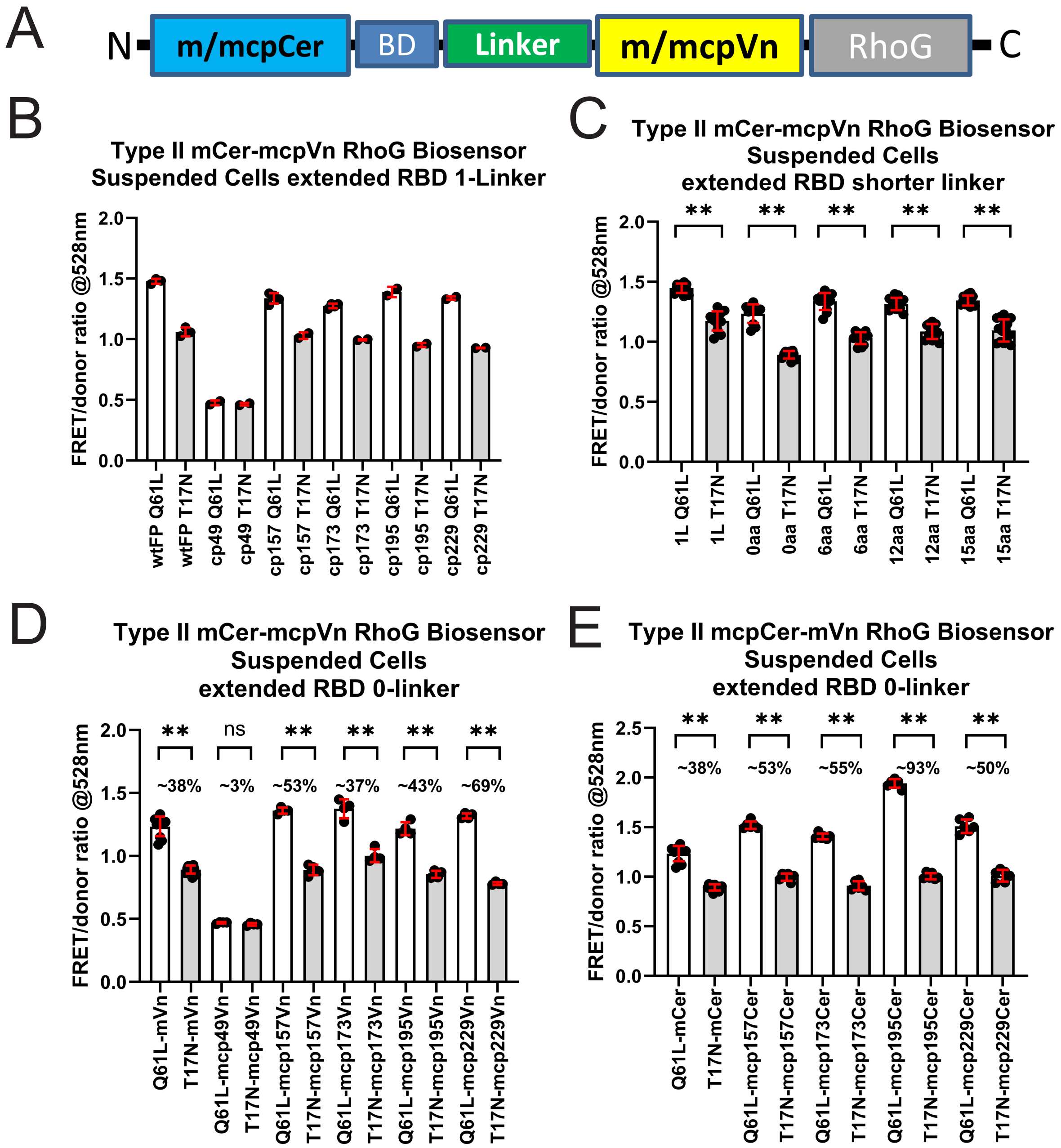


**Figure ST3: A.** Schematic drawing of Type II RhoG biosensor prototypes presented in this panel, based on mCerulean and mVenus FRET pair, with or without circular permutation. **B**. Type II RhoG biosensor with circular permutation in mVenus FRET acceptor, with extended RBD (1-330 amino acids). Detached and suspended cells measured in fluorometer. N = 2-4, shown with mean +/- SD. **C**. Type II RhoG biosensor with mCer and mVn FRET pair, extended RBD (1-330 amino acids), but with shorter linkers. Detached and suspended cells measured in fluorometer. N = 11-15, shown with mean +/- SD. Student-t test (unpaired), ** p<0.01. **D**. Type II RhoG biosensor with mCer and mcpVn FRET pair, extended RBD (1-330 amino acids), with 0-linker. Detached and suspended cells measured in fluorometer. N = 4-11, shown with mean +/- SD. Student-t test (unpaired), ** p<0.01. **E**. Type II RhoG biosensor with mcpCer and mVn FRET pair, extended RBD (1-330 amino acids), with 0-linker. Detached and suspended cells measured in fluorometer. N = 6-11, shown with mean +/- SD. Student-t test (unpaired), ** p<0.01.

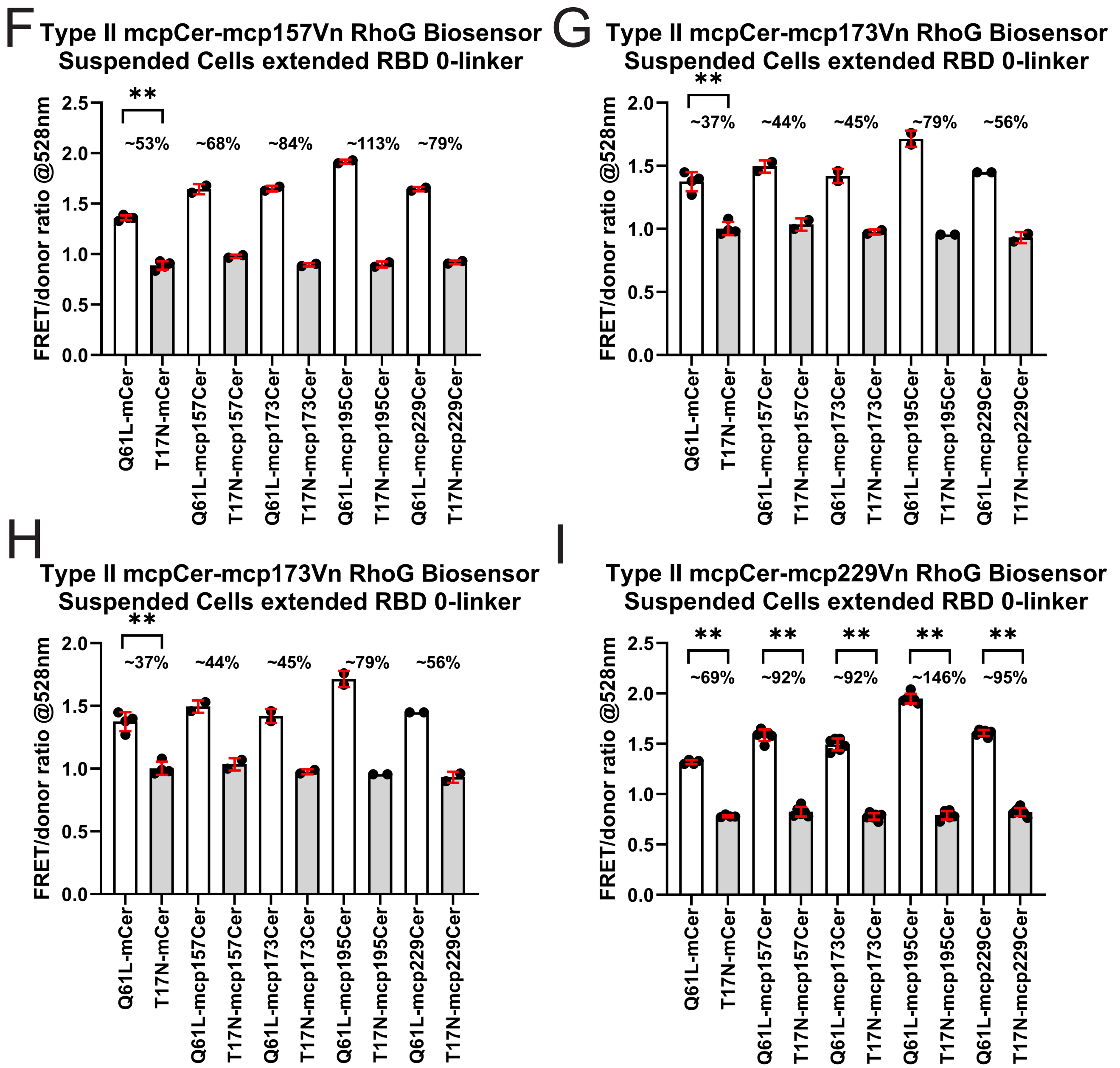


**Figure ST3 cont'd: F**. Type II RhoG biosensor with circular permutation in mCer and mcp157Vn FRET pair, extended RBD (1-330 amino acids). Detached and suspended cells measured in fluorometer. N = 2-4, shown with mean +/- SD. Student-t test (unpaired), ** $p<0.01$. **G**. Type II RhoG biosensor with circular permutation in mCer and mcp173Vn FRET pair, extended RBD (1-330 amino acids). Detached and suspended cells measured in fluorometer. N = 2-4, shown with mean +/- SD. Student-t test (unpaired), ** $p<0.01$. **H**. Type II RhoG biosensor with circular permutation in mCer and mcp195Vn FRET pair, extended RBD (1-330 amino acids). Detached and suspended cells measured in fluorometer. N = 2-4, shown with mean +/- SD. Student-t test (unpaired), ** $p<0.01$. **I**.Type II RhoG biosensor with circular permutation in mCer and mcp229Vn FRET pair, extended RBD (1-330 amino acids). Detached and suspended cells measured in fluorometer. N = 4-6, shown with mean +/- SD. Student-t test (unpaired), ** $p<0.01$.

appeared to perform better with shorter linker lengths between the FRET donor and acceptor. We therefore tested linker variants that were shorter than the original one-linker construct, which contained 17 amino acids (Fig. ST3B). Direct attachment of the two FRET moieties without an intervening linker, apart from restriction-site residues, produced the strongest response (Fig. ST3C). We therefore continued optimization using this zero-linker version of the biosensor and tested circularly permuted variants of both the mCerulean donor and mVenus acceptor. A combination of mcp195Cer as the donor and mcp229Vn as the acceptor produced the strongest overall change in FRET/donor ratio between the Q61L and T17N states, reaching approximately 146% (Fig. ST3D–I). However, the overall expression levels of these biosensors were relatively low. Although mCerulean and mVenus are bright fluorescent proteins, including after circular permutation, the raw fluorescence intensities of these constructs were approximately an order of magnitude lower than those of previous biosensors in which we had used

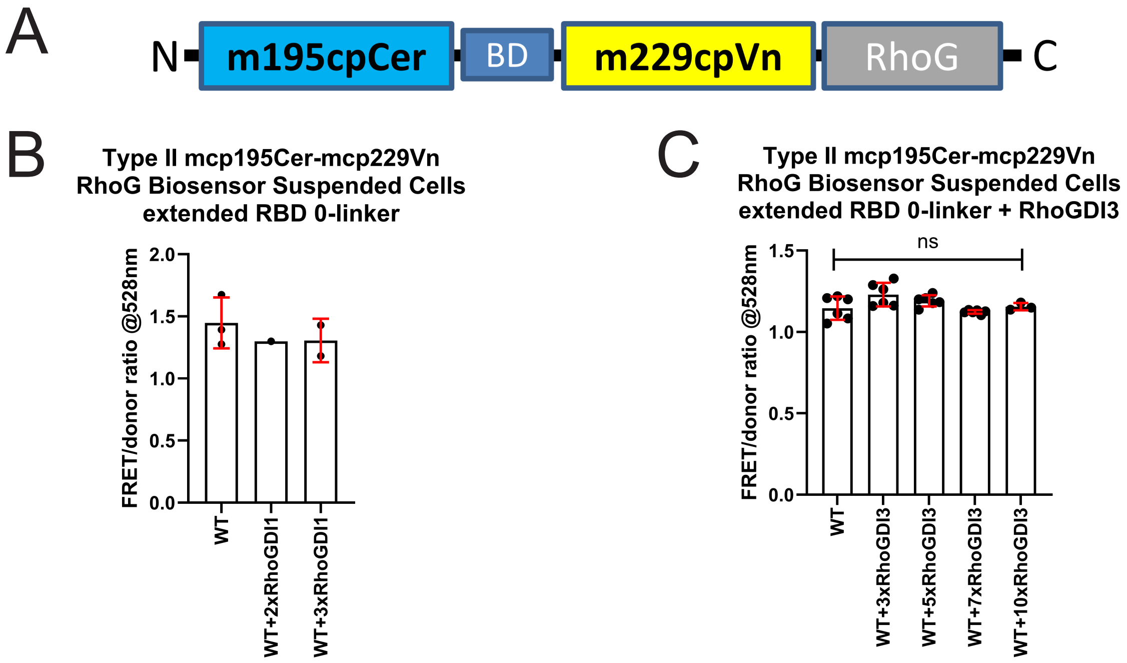


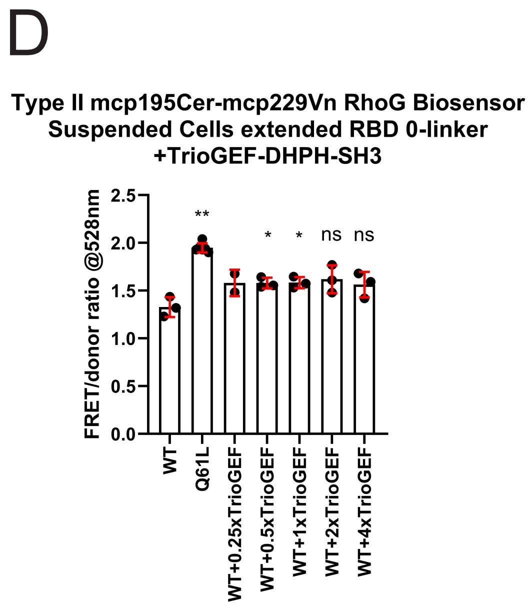


**Figure ST4: A**. Schematic drawing of Type II RhoG biosensor prototype used in this panel, based on mcp-195Cerulean and mcp229Venus FRET pair, with no internal linker segment. **B**. Type II RhoG biosensor with mcp195Cer and mcp229Vn FRET pair, extended RBD (1-330 amino acids), with 0-linker. Detached and suspended cells measured in fluorometer. N = 1-3, shown with mean +/- SD. **C**. Type II RhoG biosensor with mcp195Cer and mcp229Vn FRET pair, extended RBD (1-330 amino acids), with 0-linker. Titration of RhoGDI3 co-expression. Detached and suspended cells measured in fluorometer. N = 3-6, shown with mean +/- SD. **D**. Type II RhoG biosensor with mcp-195Cer and mcp229Vn FRET pair, extended RBD (1-330 amino acids), with 0-linker. Titration of TrioGEF-DHPH1-SH3 fragment co-expression. Detached and suspended cells measured in fluorometer. N = 2-6, shown with mean +/- SD. Panels **C** and **D**, Student-t test (unpaired), ** p<0.01, * p<0.05, ns = not significant, compared to expression of WT biosensor only.

the same circularly permuted fluorescent protein variants. This suggested that other components of the biosensor, rather than the fluorescent proteins themselves, limited biosensor expression or stability.

We next tested whether the optimized RhoG biosensor responded appropriately to co-expression of known regulators (Fig. ST4). Co-expression of RhoGDI1 at a two- to threefold excess relative to the biosensor cDNA produced no discernible change in the FRET/donor ratio compared with the WT biosensor alone (Fig. ST4B), indicating suboptimal detection of this inhibitory binding partner. Around this time, we also generated a RhoGDI3 construct and titrated it against the WT biosensor using the same co-expression assay. Similar to RhoGDI1, RhoGDI3 co-expression did not produce a significant change in the FRET/donor ratio compared with the WT biosensor alone (Fig. ST4C). The absence of a biosensor response to GDI co-expression was a strong indicator of suboptimal biosensor behavior. Cellular Rho GTPases are commonly maintained in inactive, cytoplasmic complexes with RhoGDIs (108). Therefore, a biosensor that does not detect GDI-associated inhibition may not accurately report an important regulatory transition between inactive and active RhoG states. Conversely, we tested whether the WT biosensor responded to co-expression of a known RhoG activator. We used the TrioGEF-DHPH1-SH3 fragment (92, 138), which has been reported to activate RhoG. We expected this fragment to drive the WT biosensor toward FRET/donor ratio values similar to those observed with the constitutively active Q61L mutant. However, titrated co-expression of this GEF fragment produced only a modest increase in the FRET/donor ratio of the WT biosensor (Fig. ST4D).

In parallel, we tested expression of the ELMO1 BD fragment used in this biosensor design to determine whether this domain contributed to the low cellular expression of the biosensor. The ELMO1 BD fragment comprising amino acids 1–330 was fused to GST for bacterial expression and purification. Unlike the shorter ELMO1 fragments comprising amino acids 1–115 or 1–80, the 1–330 fragment failed to express detectably in bacteria under these conditions (Fig. ST5). This result suggested that the low expression of the corresponding RhoG biosensor in cells may have been caused, at least in part, by instability or poor expression of the extended ELMO1-binding domain.

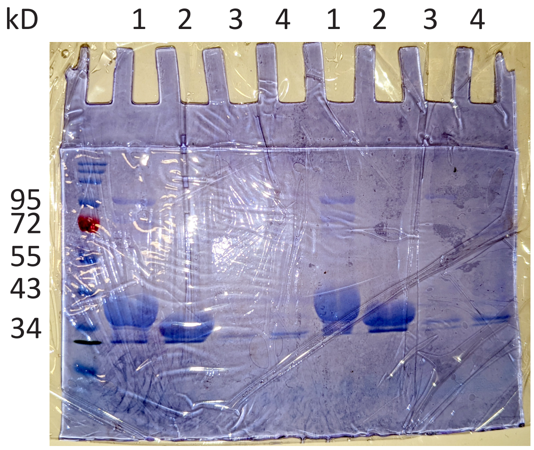


**Figure ST5**: Coomassie blue staining of an SDS-PAGE gel (duplicate loadings), showing bacterially expressed and purified GST-fusion of various ELMO1 RBD domains. Lane: 1. ELMO1 RBD 1-115 amino acids; 2. ELMO1 RBD 1-80 amino acids; 3. ELMO1 RBD 1-330 amino acids; and 4. ELMO1 RBD 1-330 amino acids containing L43A mutation.

Together, these observations indicated that, despite the large apparent difference between the Q61L and T17N mutant states, this version of the RhoG biosensor had suboptimal expression and limited responsiveness to upstream regulators. We therefore concluded that this design was not appropriate for further validation or biological analysis.

**Biosensors based on the mECFP–mCitrine FRET pair**

We next re-engineered the biosensor using the ECFP–Citrine FRET pair. We also introduced the A206K monomerizing mutation into both fluorescent proteins, because this mutation has been shown to improve off-state FRET decoupling in related biosensor designs (139). The use of mECFP as the FRET donor was intended to slightly reduce the Förster radius of the biosensor. Because mECFP has a lower quantum yield

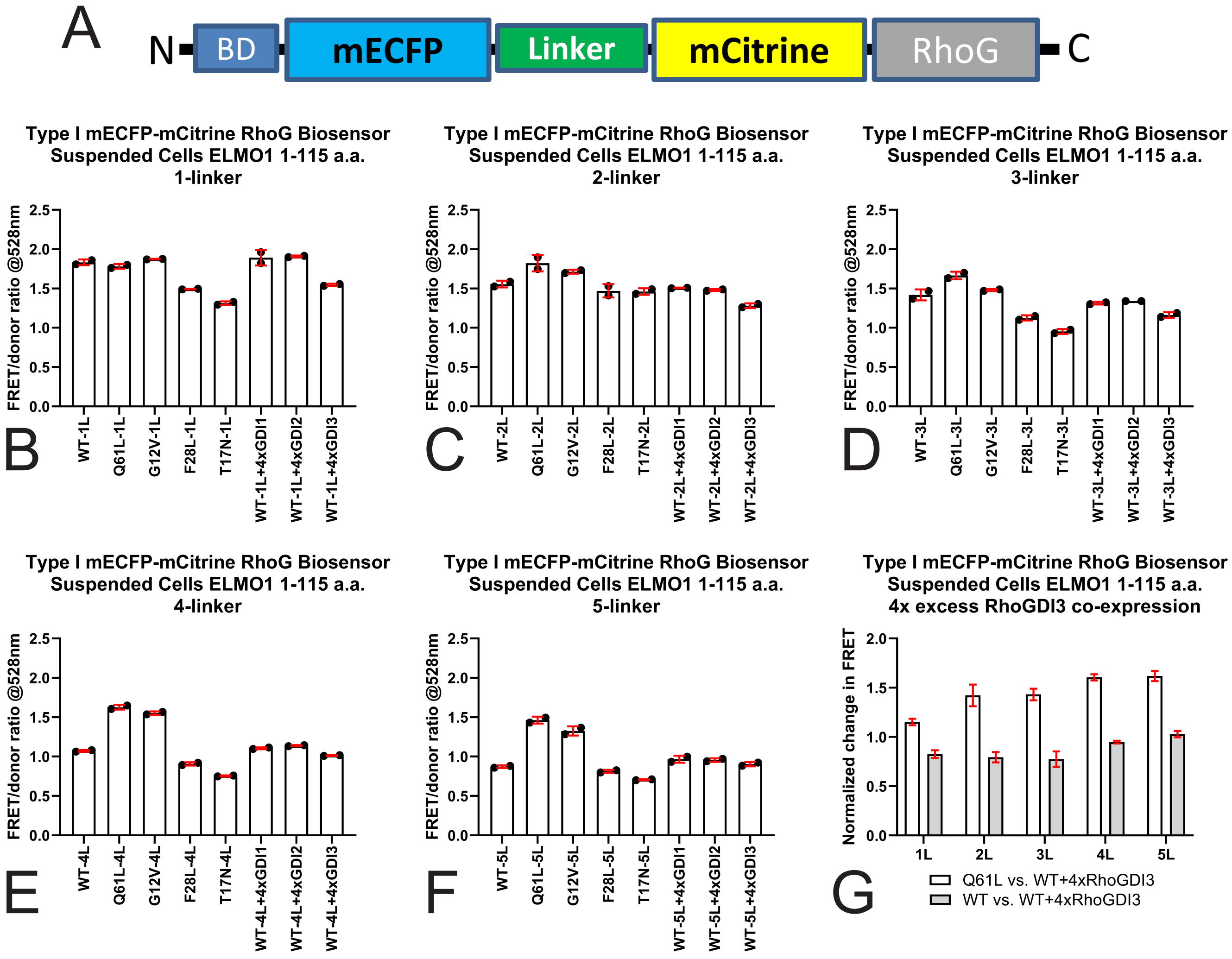


**Figure ST6: A.** Schematic drawing of Type I RhoG biosensor prototype used in this panel, based on mECFP and mCitrine FRET pair. **B.** Type I RhoG biosensor with mECFP and mCitrine FRET pair, ELMO1 RBD (1-115 amino acids) with 1-linker. Detached and suspended cells measured in fluorometer. N = 2, shown with mean +/- data range. **C.** Type I RhoG biosensor with mECFP and mCitrine FRET pair, ELMO1 RBD (1-115 amino acids) with 2-linker. Detached and suspended cells measured in fluorometer. N = 2, shown with mean +/- data range. **D.** Type I RhoG biosensor with mECFP and mCitrine FRET pair, ELMO1 RBD (1-115 amino acids) with 3-linker. Detached and suspended cells measured in fluorometer. N = 2, shown with mean +/- data range. **E.** Type I RhoG biosensor with mECFP and mCitrine FRET pair, ELMO1 RBD (1-115 amino acids) with 4-linker. Detached and suspended cells measured in fluorometer. N = 2, shown with mean +/- data range. **F.** Type I RhoG biosensor with mECFP and mCitrine FRET pair, ELMO1 RBD (1-115 amino acids) with 5-linker. Detached and suspended cells measured in fluorometer. N = 2, shown with mean +/- data range. **G.** Normalized change in FRET/donor ratio comparing 4x excess RhoGDI3 co-expression against expression of the Q61L or the WT biosensor only. Data from panels **B**-**F** were used, shown with mean +/- data range.

than mCerulean (0.41 versus 0.49) (132, 140), it should reduce the Förster radius and potentially improve discrimination between biosensor states when intramolecular movements are relatively small. Consistent with this rationale, the calculated Förster distances for the mCerulean–mVenus and mECFP–mCitrine pairs were 52.66 Å and 50.26 Å, respectively ([www.fpbase.org](http://www.fpbase.org)). Using this approach, we again generated Type I and Type II designs, each containing one to five intramolecular linker units. For these constructs, we used the shorter ELMO1 BD fragment comprising amino acids 1–115 to improve expression. In the Type I

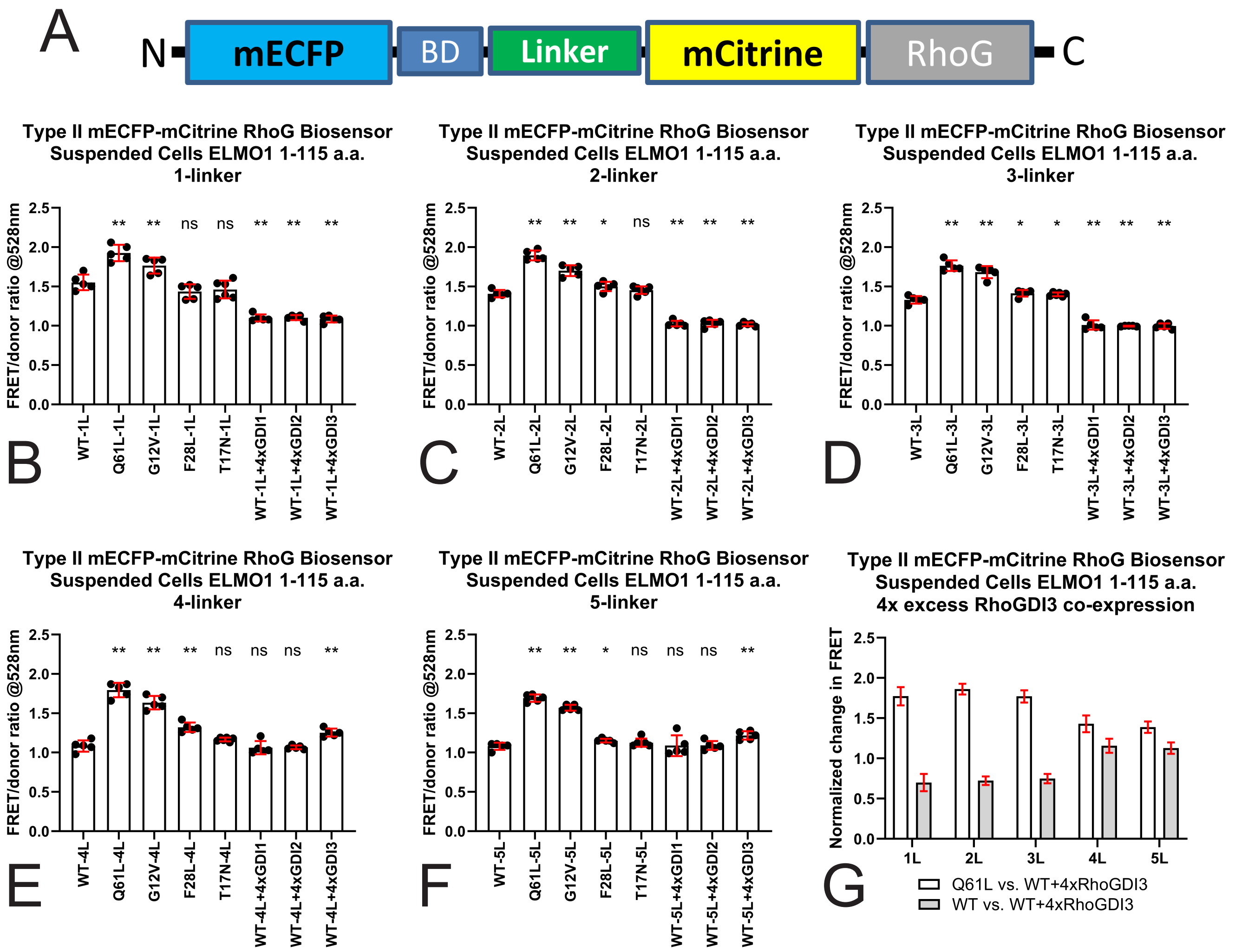


**Figure ST7: A.** Schematic drawing of Type II RhoG biosensor prototype used in this panel, based on mECFP and mCitrine FRET pair. **B**. Type II RhoG biosensor with mECFP and mCitrine FRET pair, ELMO1 RBD (1-115 amino acids) with 1-linker. Detached and suspended cells measured in fluorometer. N = 5-6, shown with mean +/- SD. **C**. Type II RhoG biosensor with mECFP and mCitrine FRET pair, ELMO1 RBD (1-115 amino acids) with 2-linker. Detached and suspended cells measured in fluorometer. N = 5-6, shown with mean +/- SD. **D**. Type II RhoG biosensor with mECFP and mCitrine FRET pair, ELMO1 RBD (1-115 amino acids) with 3-linker. Detached and suspended cells measured in fluorometer. N = 5-6, shown with mean +/- SD. **E**. Type II RhoG biosensor with mECFP and mCitrine FRET pair, ELMO1 RBD (1-115 amino acids) with 4-linker. Detached and suspended cells measured in fluorometer. N = 5-6, shown with mean +/- SD. **F**. Type II RhoG biosensor with mECFP and mCitrine FRET pair, ELMO1 RBD (1-115 amino acids) with 5-linker. Detached and suspended cells measured in fluorometer. N = 5-6, shown with mean +/- SD. **G**. Normalized change in FRET/donor ratio comparing 4x excess RhoGDI3 co-expression against expression of the Q61L or the WT biosensor only. Data from panels **B**-**F** were used, shown with mean +/- SD. Panels **B**-**F**, Student-t test (unpaired), ** $p<0.01$, * $p<0.05$, ns = not significant, compared to expression of WT biosensor only.

design, we tested WT, Q61L, G12V, F28L, and T17N RhoG variants, as well as WT biosensor co-expressed with a fourfold excess of RhoGDI1, RhoGDI2, or RhoGDI3 (Fig. ST6). The change in FRET ratio comparing the Q61L activated mutant version versus the WT with fourfold co-expression of RhoGDI3 were modest and reached a maximum of 1.62x change in the five-linker version of the biosensor (Fig. ST6G). RhoGDI3-

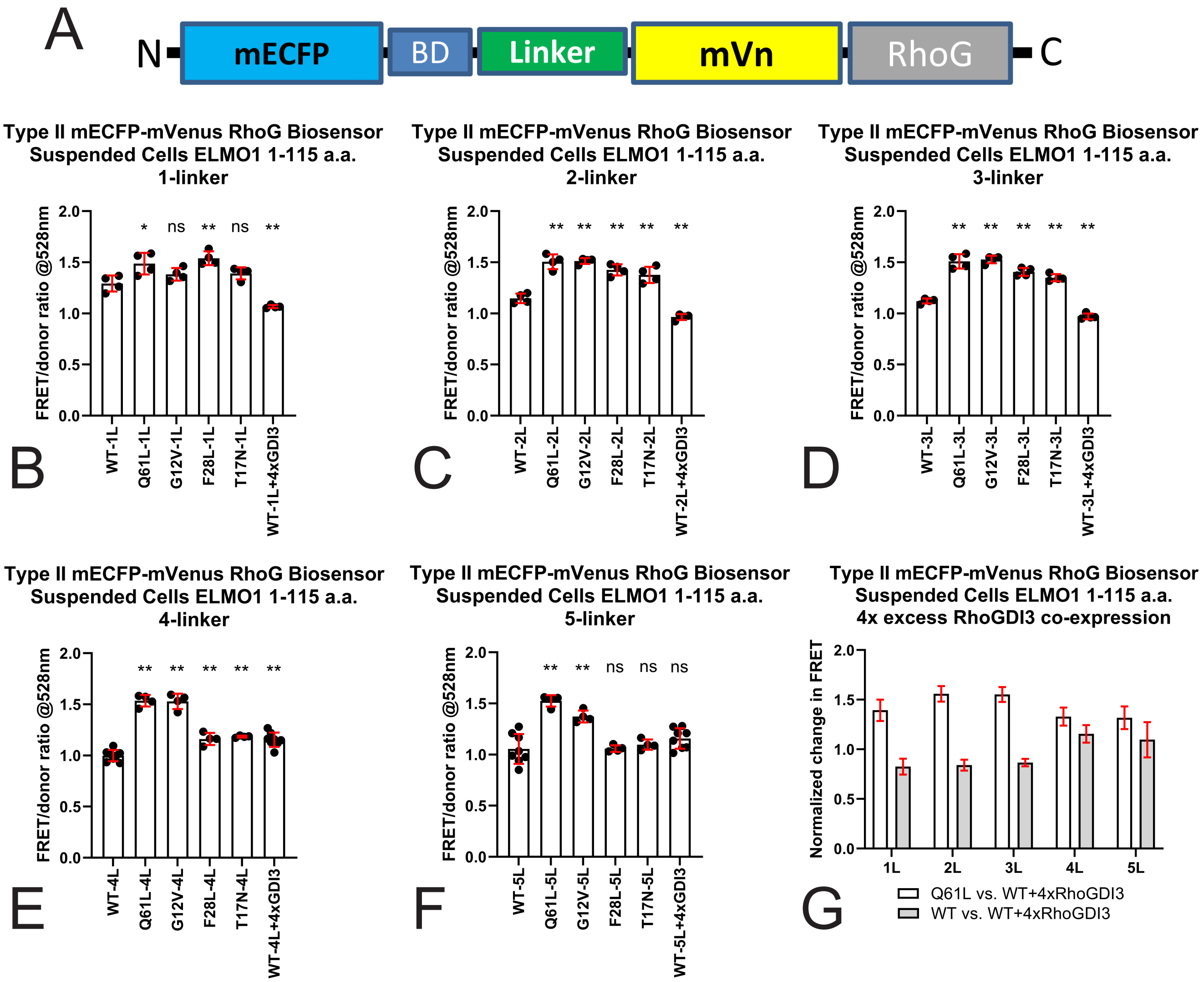


**Figure ST8:** **A.** Schematic drawing of Type II RhoG biosensor prototype used in this panel, based on mECFP and mVenus FRET pair. **B**. Type II RhoG biosensor with mECFP and mVenus FRET pair, ELMO1 RBD (1-115 amino acids) with 1-linker. Detached and suspended cells measured in fluorometer. N = 4, shown with mean +/- SD. **C**. Type II RhoG biosensor with mECFP and mVenus FRET pair, ELMO1 RBD (1-115 amino acids) with 2-linker. Detached and suspended cells measured in fluorometer. N = 4, shown with mean +/- SD. **D**. Type II RhoG biosensor with mECFP and mVenus FRET pair, ELMO1 RBD (1-115 amino acids) with 3-linker. Detached and suspended cells measured in fluorometer. N = 4, shown with mean +/- SD. **E**. Type II RhoG biosensor with mECFP and mVenus FRET pair, ELMO1 RBD (1-115 amino acids) with 4-linker. Detached and suspended cells measured in fluorometer. N = 4-8, shown with mean +/- SD. **F**. Type II RhoG biosensor with mECFP and mVenus FRET pair, ELMO1 RBD (1-115 amino acids) with 5-linker. Detached and suspended cells measured in fluorometer. N = 4-8, shown with mean +/- SD. All panels, Student-t test (unpaired), ** p<0.01, * p<0.05, ns = not significant, compared to expression of WT biosensor only. **G**. Normalized change in FRET/donor ratio comparing 4x excess RhoGDI3 co-expression against expression of the Q61L or the WT biosensor only. Data from panels **B**-**F** were used, shown with mean +/- SD.

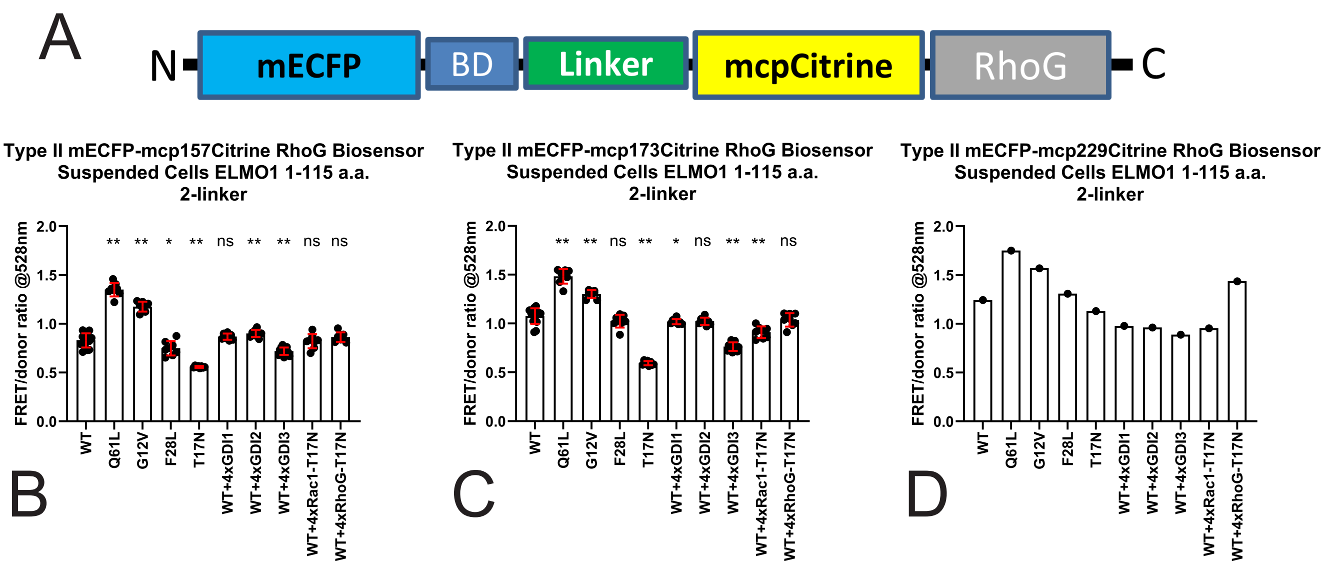


**Figure ST9: A.** Schematic drawing of Type II RhoG biosensor prototype used in this panel, based on mECFP and mcpCitrine FRET pair. **B**. Type II RhoG biosensor with mECFP and mcp157Citrine FRET pair, ELMO1 RBD (1-115 amino acids) with 2-linker. Detached and suspended cells measured in fluorometer. N = 8-14, shown with mean +/- SD. **C**. Type II RhoG biosensor with mECFP and mcp173Citrine FRET pair, ELMO1 RBD (1-115 amino acids) with 2-linker. Detached and suspended cells measured in fluorometer. N = 8-14, shown with mean +/- SD. **D**. Type II RhoG biosensor with mECFP and mcp229Citrine FRET pair, ELMO1 RBD (1-115 amino acids) with 2-linker. Detached and suspended cells measured in fluorometer. N = 1. In panels **B** and **C**, Student-t test (unpaired), ** p<0.01, * p<0.05, ns = not significant, compared to expression of WT biosensor only.

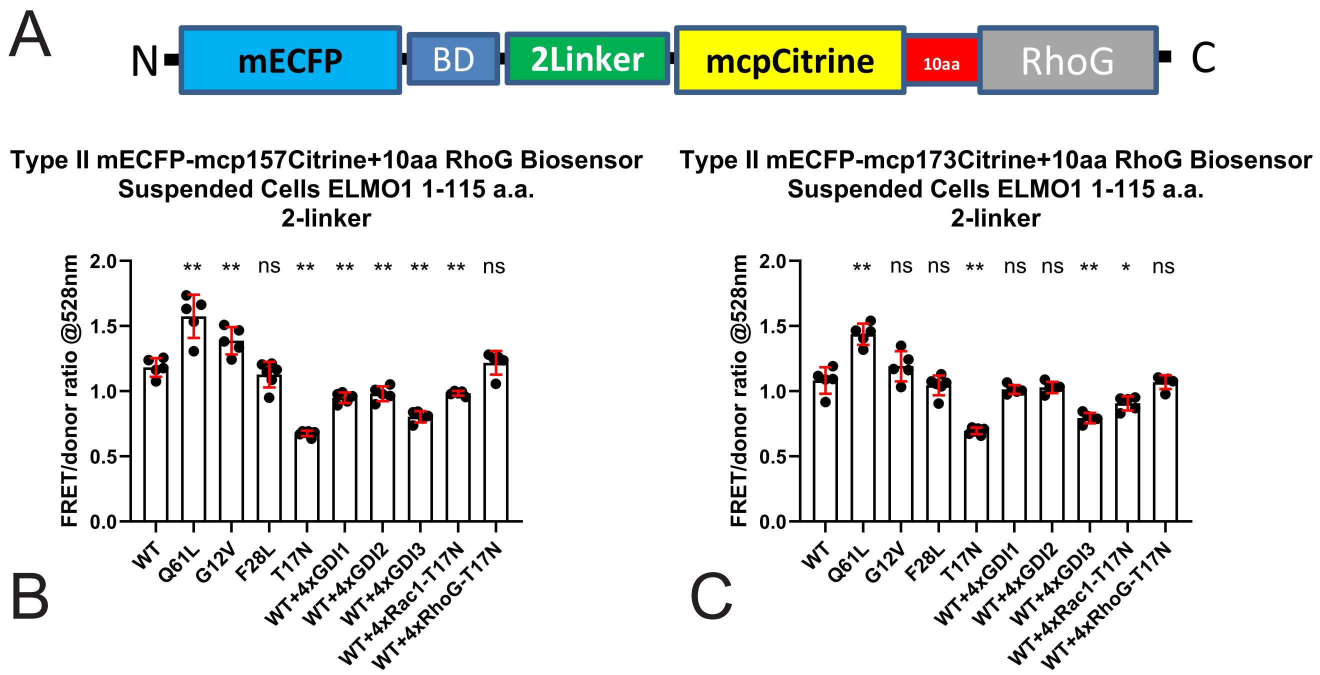


**Figure ST10: A**. Schematic drawing of Type II RhoG biosensor prototype, utilizing mECFP and mcpCitrine, with 2 linker internal linker structure plus additional 10 amino acids being added in between mcpCitrine and RhoG. **B**. Type II RhoG biosensor with mECFP and mcp157Citrine FRET pair, ELMO1 RBD (1-115 amino acids) with 2-linker, with 10 amino acids terminal extension on mcpCitrine. Detached and suspended cells measured in fluorometer. N = 5-6, shown with mean +/- SD. **C**. Type II RhoG biosensor with mECFP and mcp173Citrine FRET pair, ELMO1 RBD (1-115 amino acids) with 2-linker, with 10 amino acids terminal extension on mcpCitrine. Detached and suspended cells measured in fluorometer. N = 8-14, shown with mean +/- SD. In panels **B** and **C**, Student-t test (unpaired), ** p<0.01, * p<0.05, ns = not significant, compared to expression of WT biosensor only.

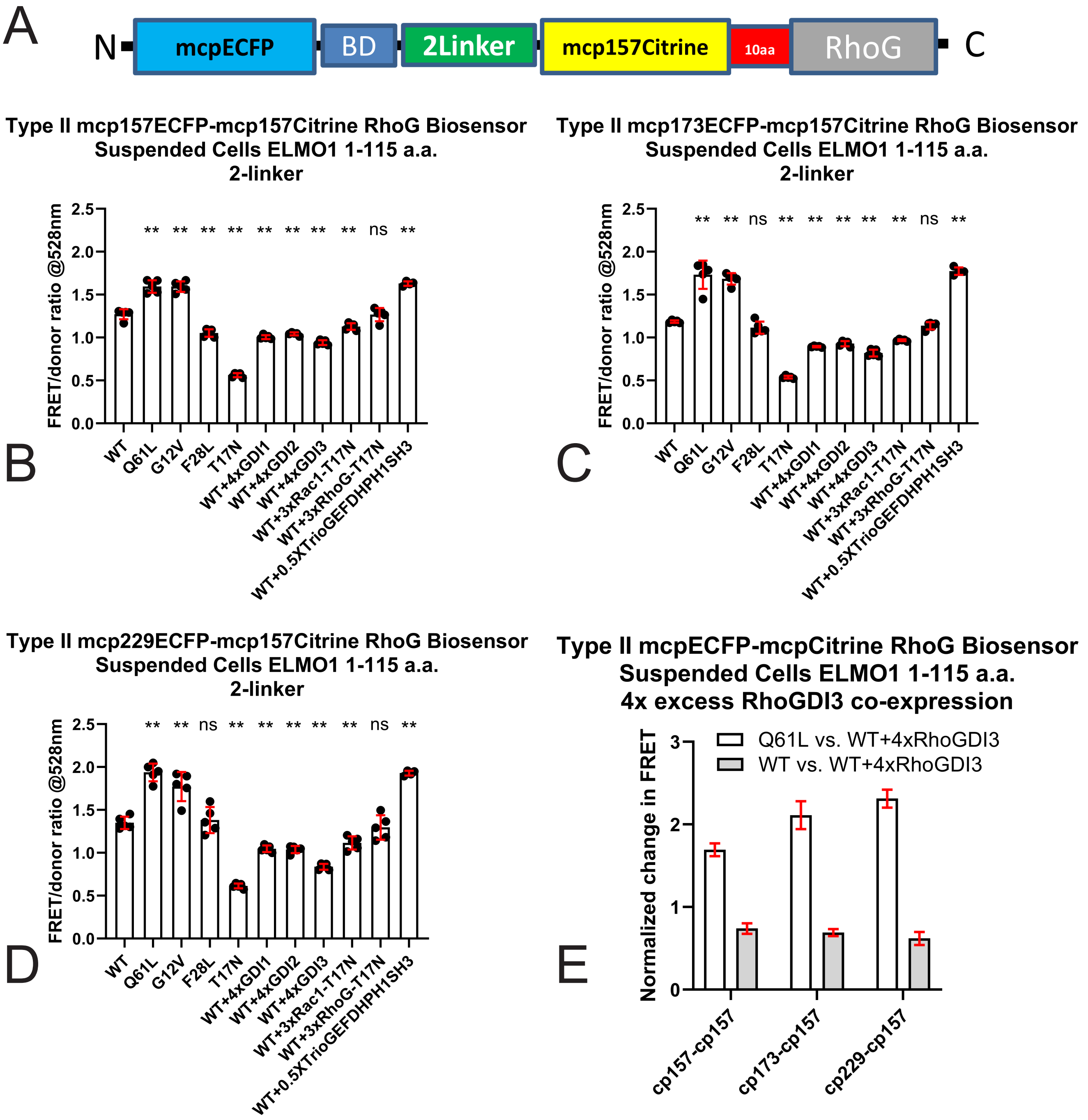


**Figure ST11: A**. Schematic drawing of Type II RhoG biosensor prototype, utilizing mcpECFP and mcp-157Citrine, with 2 internal linker structure plus additional 10 amino acids being added in between mcp-157Citrine and RhoG. **B**. Type II RhoG biosensor with m157ECFP and mcp157Citrine FRET pair, ELMO1 RBD (1-115 amino acids) with 2-linker, with 10 amino acids terminal extension on mcp157Citrine. Detached and suspended cells measured in fluorometer. N = 4-5, shown with mean +/- SD. **C**. Type II RhoG biosensor with mcp173ECFP and mcp157Citrine FRET pair, ELMO1 RBD (1-115 amino acids) with 2-linker, with 10 amino acids terminal extension on mcp157Citrine. Detached and suspended cells measured in fluorometer. N = 4-5, shown with mean +/- SD. **D**. Type II RhoG biosensor with mcp229ECFP and mcp157Citrine FRET pair, ELMO1 RBD (1-115 amino acids) with 2-linker, with 10 amino acids terminal extension on mcp157Citrine. Detached and suspended cells measured in fluorometer. N = 4-5, shown with mean +/- SD. In panels **B**, **C**, and **D**, Student-t test (unpaired), ** $p<0.01$, * $p<0.05$, ns = not significant, compared to expression of WT biosensor only. **E**. Normalized change in FRET/donor ratio comparing 4x excess RhoGDI3 co-expression against expression of the Q61L or the WT biosensor only. Data from panels **B**-**D** were used, shown with mean +/- SD.

dependent reduction in FRET/donor ratio was observed up to the three-linker version. However, the four- and five-linker versions showed little or no response to GDI co-expression, suggesting that these longer linker configurations produced suboptimal structural orientations that did not appropriately report the GDI-associated inhibitory state (Fig. ST6G).

We then turned to the Type II design. In this configuration, we tested WT, Q61L, G12V, F28L, and T17N RhoG variants, as well as WT biosensor co-expressed with a fourfold excess of RhoGDI1, RhoGDI2, or RhoGDI3 (Fig. ST7). Based on the response trends, the two-linker version produced the largest difference in FRET/donor ratio between the Q61L mutant and the WT biosensor co-expressed with fourfold excess RhoGDI3, with an approximately 1.86-fold change (Fig. ST7G). In contrast, longer linkers produced elevated FRET/donor ratios in the WT biosensor co-expressed with excess RhoGDI3 (Fig. ST7G), which was considered suboptimal. We also tested the same Type II configurations after replacing the FRET acceptor with mVenus (Fig. ST8). In this case, the two-linker version again produced the greatest difference in FRET/donor ratio between Q61L and WT plus fourfold excess RhoGDI3. However, the overall change was smaller than that observed with mCitrine, reaching 1.57-fold (Fig. ST8G).

We next tested circular permutation of mCitrine in the Type II, two-linker biosensor (Fig. ST9). For mCitrine, the cp49 and cp195 variants did not fluoresce strongly, unlike the corresponding Venus variants, indicating suboptimal folding or maturation. We therefore tested mcp157, mcp173, and mcp229 versions of mCitrine as the acceptor while maintaining wild-type mECFP as the donor. For this analysis, we tested WT, Q61L, G12V, F28L, and T17N RhoG variants in the biosensor. We also tested the WT RhoG biosensor with fourfold excess RhoGDI1, RhoGDI2, or RhoGDI3; threefold excess dominant-negative Rac1 T17N or RhoG T17N; or 0.5-fold co-expression of an activated TrioGEF-DHPH1-SH3 fragment known to activate RhoG. This analysis identified the cp157 and cp173 acceptor variants as having good dynamic range and generally expected behavior across the tested conditions. The dynamic range, measured by comparing Q61L with WT plus fourfold excess RhoGDI3, was 1.88-fold for cp157 and 1.94-fold for cp173. Although cp229 showed the largest dynamic range by this metric, reaching 1.97-fold, co-expression of dominant-negative RhoG T17N unexpectedly elevated the FRET/donor ratio. We therefore removed this version from further consideration.

Next, we inserted a flexible 10-amino-acid linker between the FRET acceptor mcpCitrine and full-length RhoG (Fig. ST10A). Similar modifications in previous biosensor designs stabilized the FRET response of dominant-negative biosensor variants, likely by providing additional conformational flexibility during GTPase–GEF interactions that could otherwise spuriously alter FRET (141). With this modification, we observed that for mcp157Citrine, the ratio of T17N biosensor changed from 0.55 to 0.67 after linker insertion, and for mcp173Citrine, the ratio changed from 0.59 to 0.70. We applied this modification to both the cp157 and mcp173 versions and found that the dynamic range now favored cp157 over cp173, with 1.96-fold and 1.81-fold changes, respectively, when comparing Q61L with WT plus fourfold excess RhoGDI3 (Fig. ST10B, C). In addition, general behaviors of the mutants of biosensors, as well as responses to RhoGDI co-expressions were generally more desirable for cp157 version. We therefore selected the cp157 version for the next stage of optimization.

Finally, we tested circularly permuted versions of the mECFP donor in the Type II, two-linker biosensor containing mcp157Citrine and the 10-amino-acid linker extension between the FRET acceptor and full-length RhoG (Fig. ST11). We tested cp157, cp173, and cp229 variants of mECFP as the donor. Fluorometric measurements showed 1.69-, 2.11-, and 2.31-fold changes, respectively, when comparing the Q61L biosensor with the WT biosensor co-expressed with fourfold excess RhoGDI3 (Fig. ST11E). Importantly, co-expression of an activated TrioGEF fragment with the WT versions of these biosensors strongly increased the FRET/donor ratio to levels comparable to those observed with Q61L. This indicated that the WT biosensors retained the ability to be fully activated by a competent upstream GEF. Based on these results, we selected the cp229 version of mECFP as the FRET donor and concluded the engineering and optimization of the RhoG biosensor.
