## Supplementary figures and images for "A genetically encoded RhoG FRET biosensor reveals spatially compartmentalized RhoG–Rac1 signaling during cell protrusion"

### Supplemental Figure1

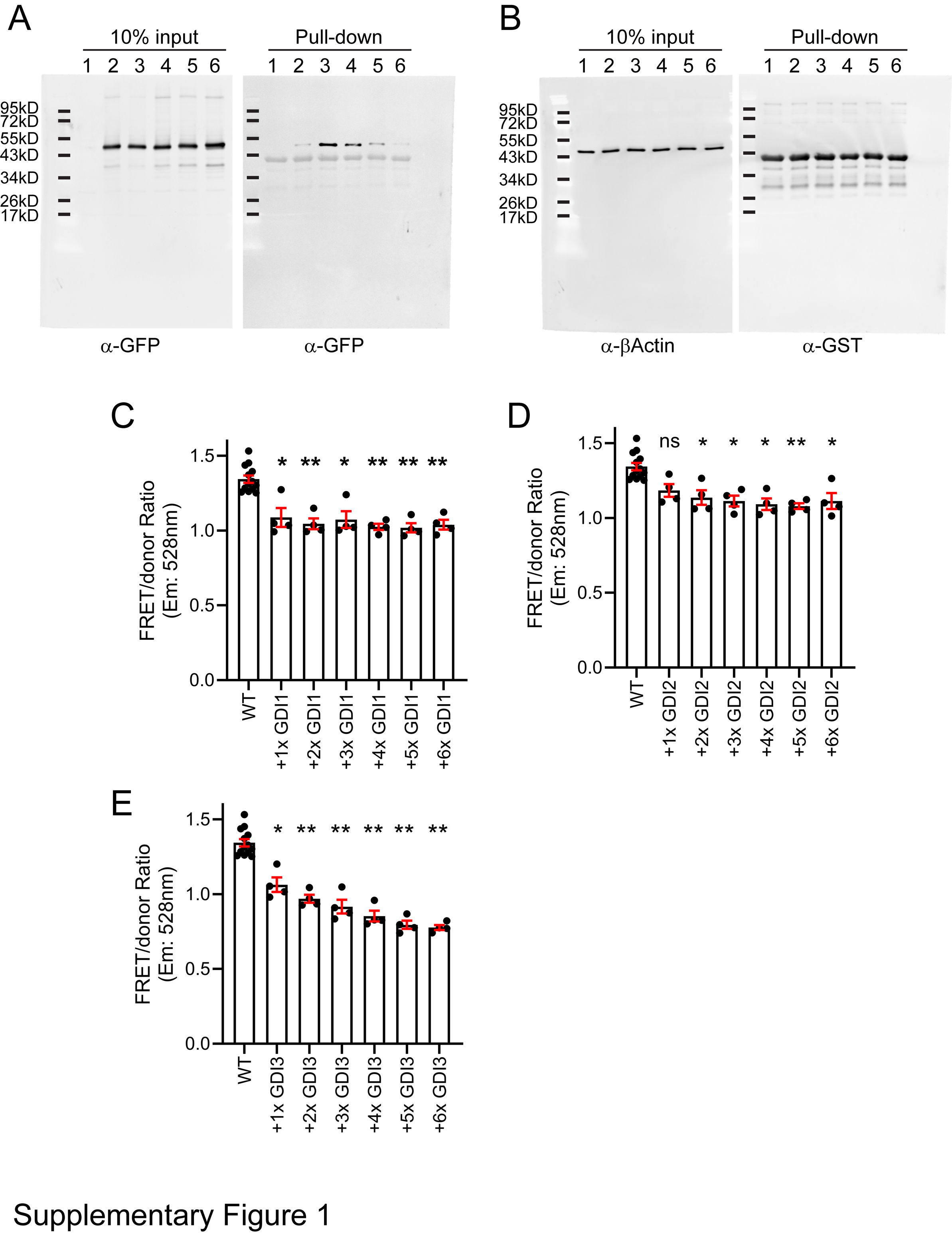

### Supplemental Figure2

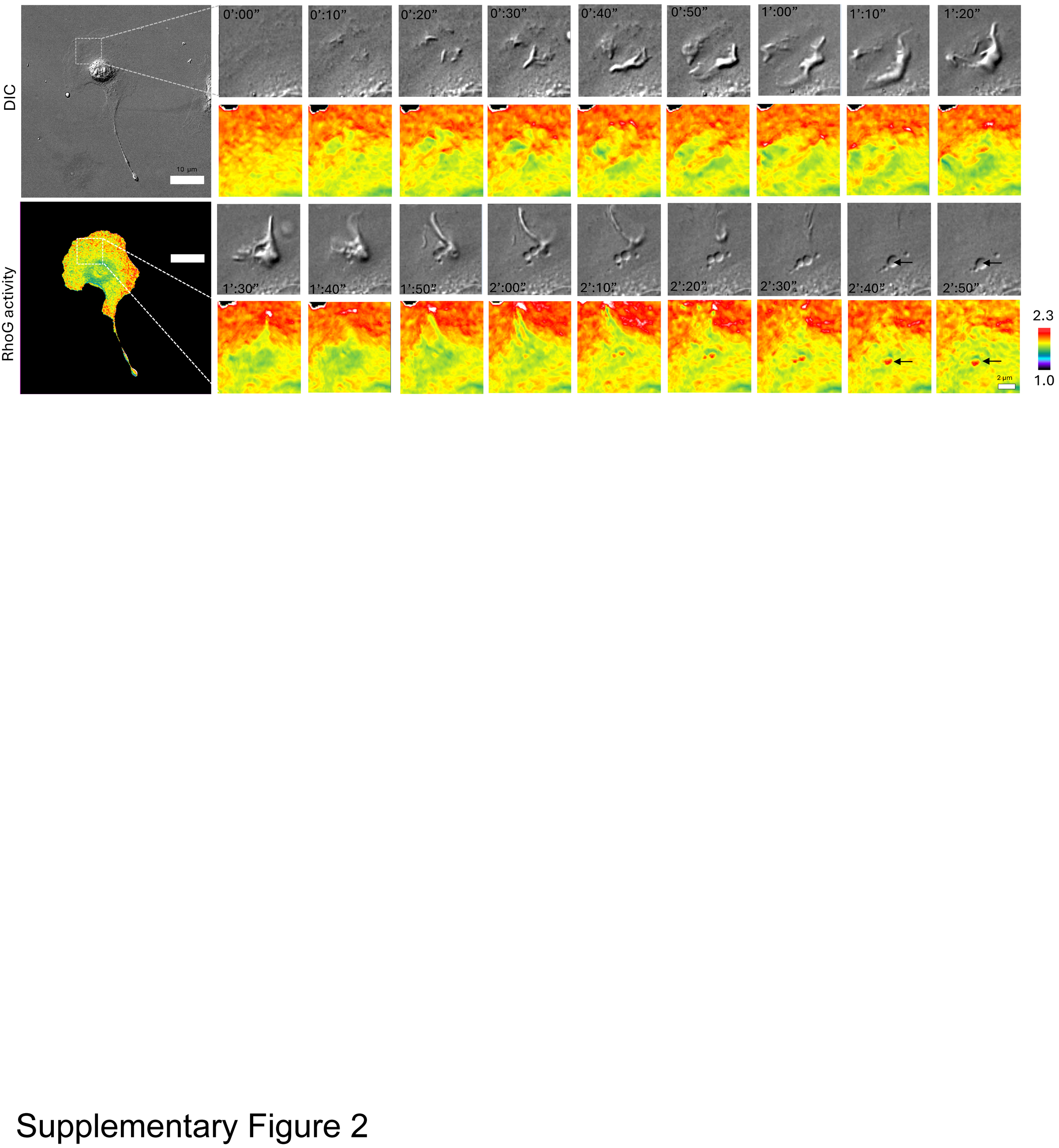

### Supplemental Figure3

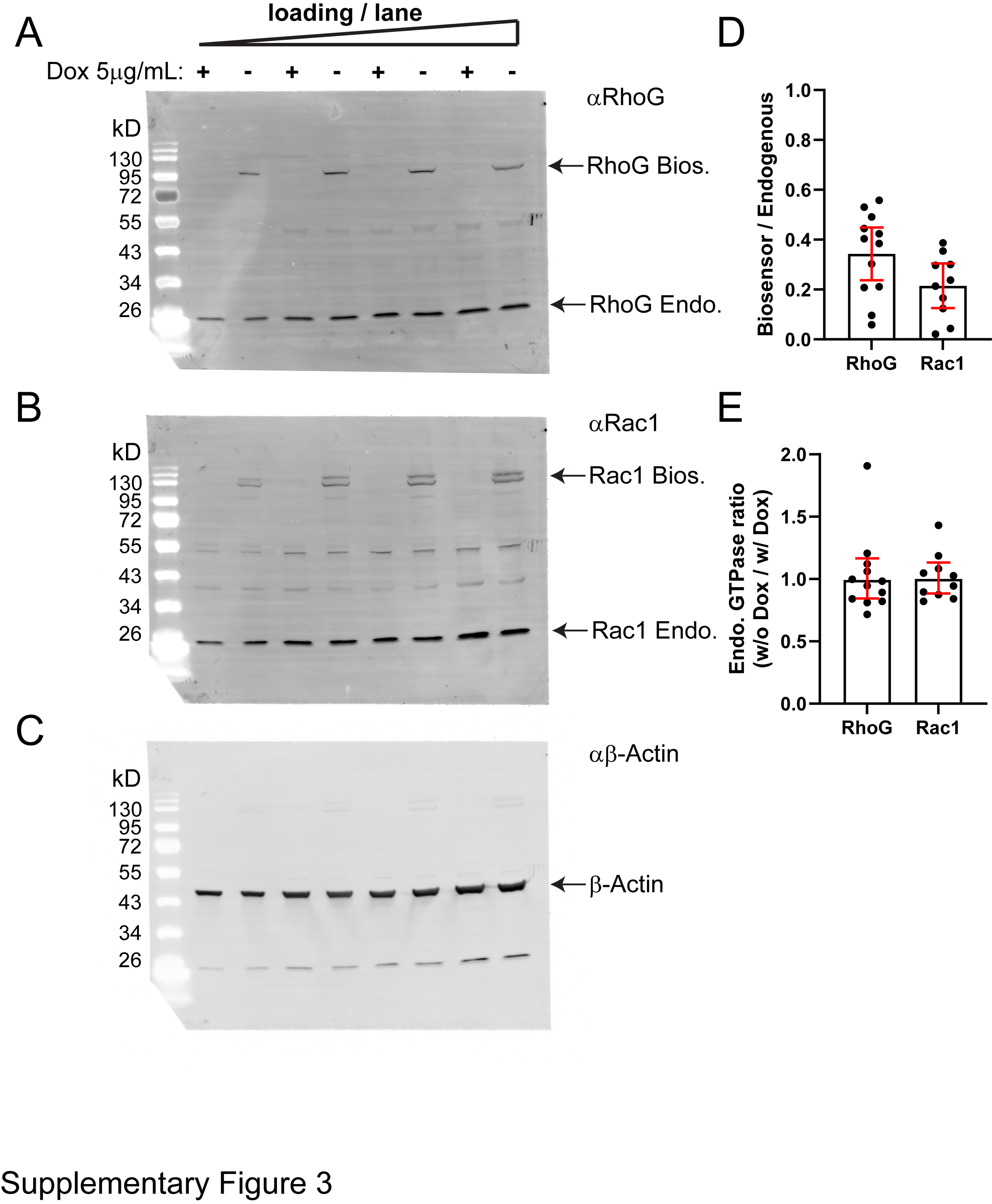

### Supplemental Figure4

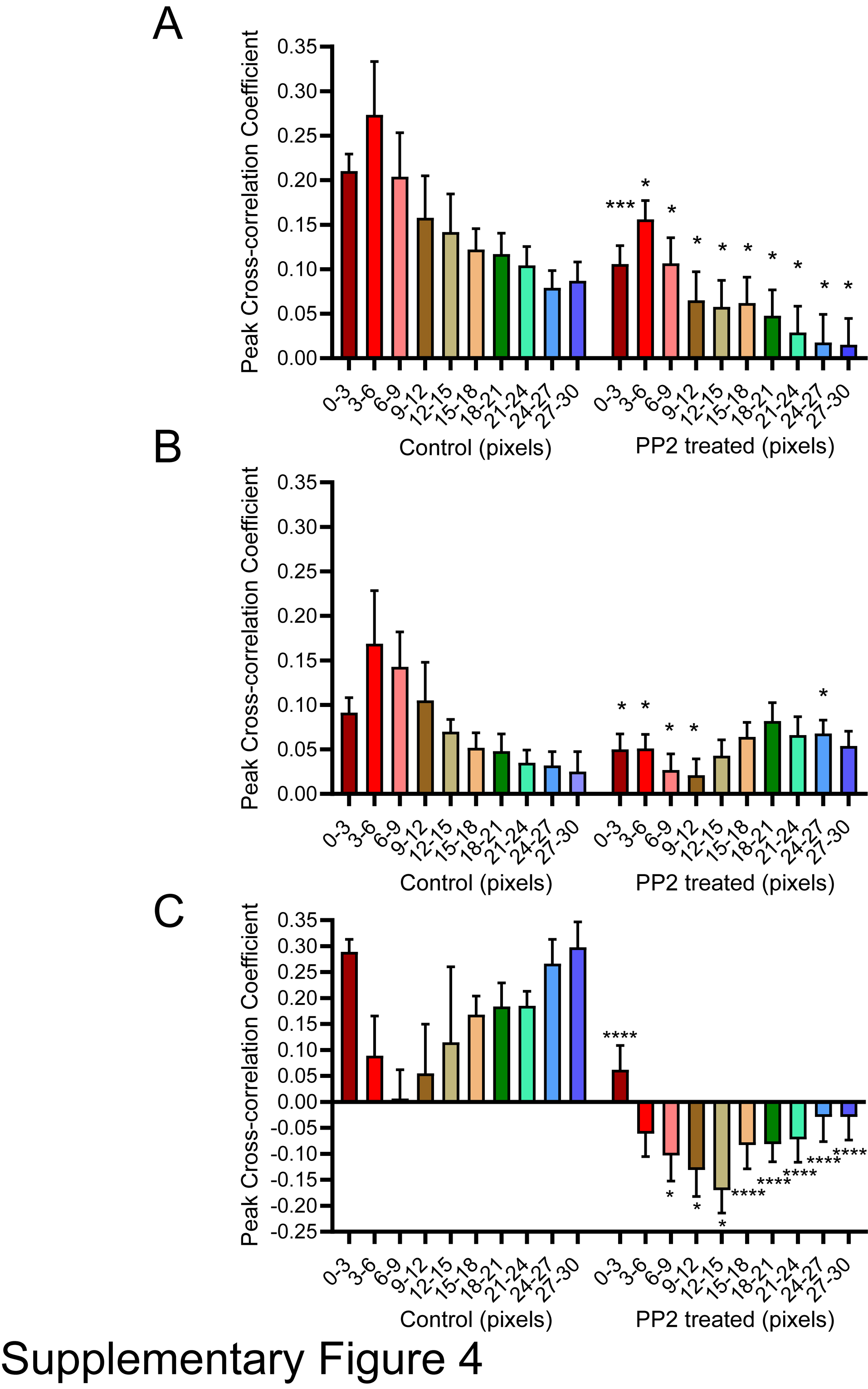

### Supplemental Figure5

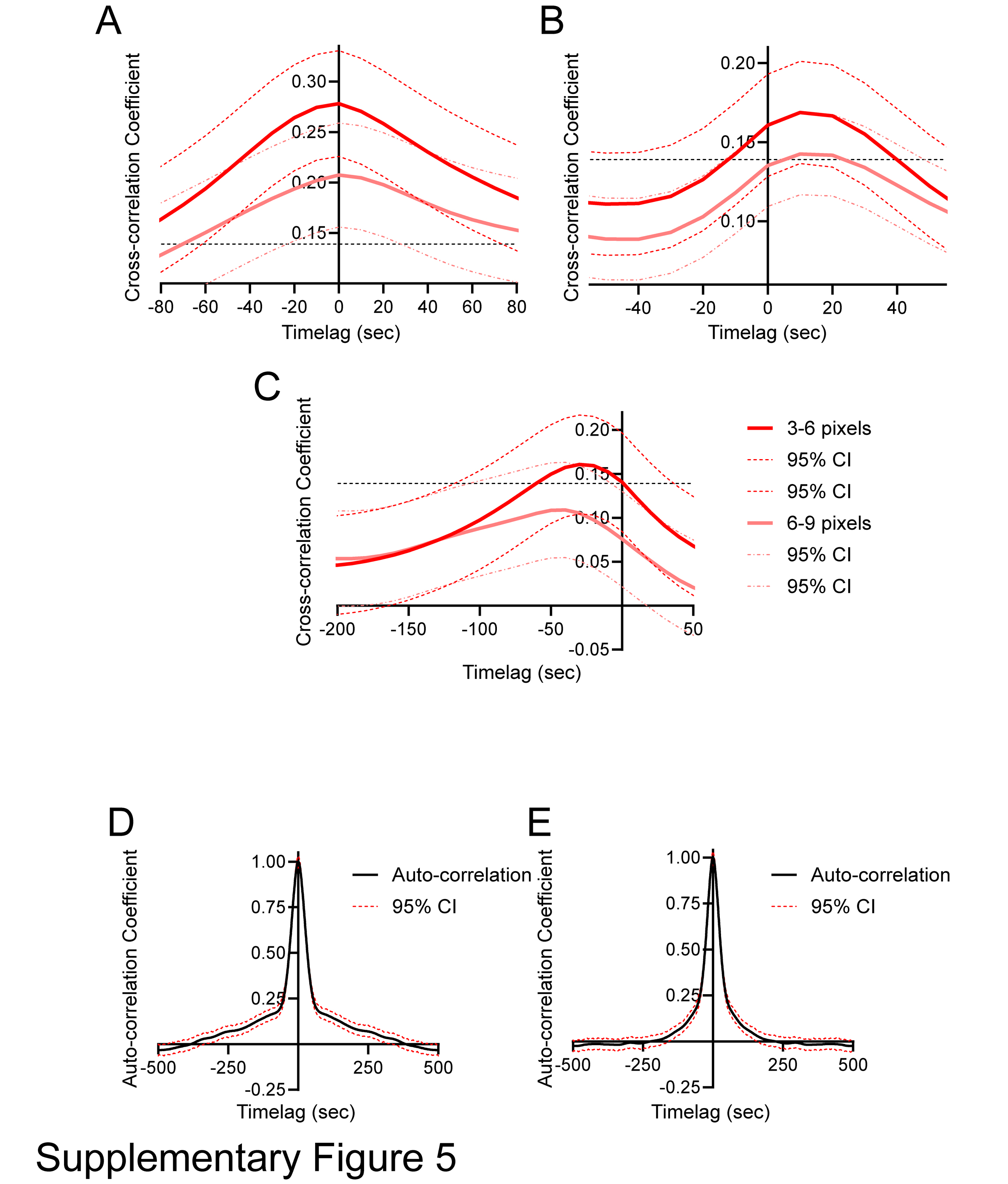
