## SupplementaryData for "A genetically encoded RhoG FRET biosensor reveals spatially compartmentalized RhoG–Rac1 signaling during cell protrusion"

Monomeric cp229ECFP-SMv1:

ATGGGCATTACCCTGGGAATGGATGAACTGTATAAAGGCGGCAGCGGCGGCATGGTGTCCAAAGGAGAAGAACTGTTTACAGGAGTGGTGCCTATTCTGGTGGAACTGGATGGAGATGTGAATGGACATCGCTTTTCCGTGAGCGGAGAAGGAGAAGGAGACGCTACATATGGAAAACTGACACTGAAGTTTATTTGTACAACAGGAAAACTGCCTGTGCCTTGGCCTACACTGGTGACCACACTGACATGGGGAGTCCAGTGTTTTAGCAGGTATCCTGATCATATGAAACAGCATGATTTCTTTAAAAGCGCTATGCCTGAGGGATATGTGCAGGAAAGGACAATTTTCTTTAAAGATGATGGAAATTATAAAACAAGGGCTGAAGTGAAATTTGAAGGAGATACACTGGTGAATAGGATTGAACTGAAAGGAATTGATTTTAAAGAAGATGGAAATATTCTGGGACATAAACTGGAATATAATTACATTAGCCATAATGTGTACATTACAGCTGATAAACAGAAAAATGGAATTAAGGCTCACTTTAAAATTAGGCATAATATTGAAGATGGAAGCGTGCAGCTGGCTGATCATTATCAGCAGAATACACCTATTGGAGATGGACCTGTGCTCCTGCCTGATAATCATTATCTGTCCACACAGAGCAAGCTGTCCAAAGATCCTAATGAAAAAAGGGACCATATGGTGCTGCTGGAATTTGTGACAGCTGCCGGA

Linker:

GGATCC

ELMO1 binding domain (1-115a.a.):

ATGCCGCCGCCTTCGGACATCGTGAAGGTGGCCATAGAATGGCCTGGCGCCTACCCTAAACTCATGGAGATTGACCAGAAAAAGCCACTGTCTGCCATTATAAAGGAAGTCTGTGATGGGTGGTCTCTTGCCAACCATGAATATTTTGCACTGCAGCACGCTGATAGTTCAAACTTCTACATCACAGAAAAGAATCGCAACGAAATAAAAAATGGCACGATCCTTCGATTAACCACATCTCCAGCTCAGAATGCCCAGCAGCTCCATGAACGAATCCAGTCCTCGTCTATGGATGCCAAGCTGGAAGCCCTGAAGGACTTGGCCAGCCTCTCCCGGGATGTTACC

Flexible linker:

GGAAGCTTAACTTCTGGTTCTGGTAAACCTGGTTCTGGTGAAGGTTCTACTAAAGGTGGATCTACTTCTGGTTCTGGTAAACCTGGTTCTGGTGAAGGTTCTACTAAAGGTGGATCTGCGGCCGCT

Monomeric cp157Citrine:

ATGCAGAAGAACGGCATCAAGGTGAACTTCAAGATCCGCCACAACATCGAGGACGGCAGCGTGCAGCTCGCCGACCACTACCAGCAGAACACCCCCATCGGCGACGGCCCCGTGCTGCTGCCCGACAACCACTACCTGAGCTACCAGTCCAAGCTGAGCAAAGACCCCAACGAGAAGCGCGATCACATGGTCCTGCTGGAGTTCGTGACCGCCGCCGGGATCACTCTCGGCATGGACGAGCTGTACAAGGGTGGCAGCGGTGGCATGGTGAGCAAGGGCGAGGAGCTGTTCACCGGGGTGGTGCCCATCCTGGTCGAGCTGGACGGCGACGTAAACGGCCACAAGTTCAGCGTGTCCGGCGAGGGCGAGGGCGATGCCACCTACGGCAAGCTGACCCTGAAGTTCATCTGCACCACCGGCAAGCTGCCCGTGCCCTGGCCCACCCTCGTGACCACCTTCGGCTACGGCCTGATGTGCTTCGCCCGCTACCCCGACCACATGAAGCAGCACGACTTCTTCAAGTCCGCCATGCCCGAAGGCTACGTCCAGGAGCGCACCATCTTCTTCAAGGACGACGGCAACTACAAGACCCGCGCCGAGGTGAAGTTCGAGGGCGACACCCTGGTGAACCGCATCGAGCTGAAGGGCATCGACTTCAAGGAGGACGGCAACATCCTGGGGCACAAGCTGGAGTACAACTACAACAGCCACAACGTCTATATCATGGCCGACAAG

Linker:

GGAAGCGGCAGCGGGTCTAGCGGAGGCAGCGAATTC

RhoG WT:

ATGCAGAGCATCAAGTGCGTGGTGGTGGGTGATGGGGCTGTGGGCAAGACGTGCCTGCTCATCTGCTACACAACTAACGCTTTCCCCAAAGAGTACATCCCCACCGTGTTCGACAATTACAGCGCGCAGAGCGCAGTTGACGGGCGCACAGTGAACCTGAACCTGTGGGACACTGCGGGCCAGGAGGAGTATGACCGCCTCCGTACACTCTCCTACCCTCAGACCAACGTTTTCGTCATCTGTTTCTCCATTGCCAGTCCGCCGTCCTATGAGAACGTGCGGCACAAGTGGCATCCAGAGGTGTGCCACCACTGCCCTGATGTGCCCATCCTGCTGGTGGGCACCAAGAAGGACCTGAGAGCCCAGCCTGACACCCTACGGCGCCTCAAGGAGCAGGGCCAGGCGCCCATCACACCGCAGCAGGGCCAGGCACTGGCCAAGCAGATCCACGCTGTGCGCTACCTCGAATGCTCAGCCCTGCAACAGGATGGTGTCAAGGAAGTGCTCGCCGAGGCTGTCCGGGCTGTGCTCAACCCCACGCCGATCAAGCGTGGGCGGTCCTGCATCCTCTTGTGA
